# Natural strain variation reveals diverse biofilm regulation in squid-colonizing *Vibrio fischeri*

**DOI:** 10.1101/471292

**Authors:** Ella R. Rotman, Katherine M. Bultman, John F. Brooks, Mattias C. Gyllborg, Hector L. Burgos, Michael S. Wollenberg, Mark J. Mandel

## Abstract

The mutualistic symbiont *Vibrio fischeri* builds a symbiotic biofilm during colonization of squid hosts. Regulation of the exopolysaccharide component, termed Syp, has been examined in strain ES114, where production is controlled by a phosphorelay that includes the inner membrane hybrid histidine kinase RscS. Most strains that lack RscS or encode divergent RscS proteins cannot colonize a squid host unless RscS from a squid symbiont is heterologously expressed. In this study, we examine *V. fischeri* isolates worldwide to understand the landscape of biofilm regulation during beneficial colonization. We provide a detailed study of three distinct evolutionary groups of *V. fischeri* and find that while the RscS-Syp biofilm pathway is required in one of the groups, two other groups of squid symbionts require Syp independent of RscS. Mediterranean squid symbionts, including *V. fischeri* SR5, colonize without an RscS homolog encoded in their genome. Additionally, Group A *V. fischeri* strains, which form a tightly-related clade of Hawaii isolates, have a frameshift in *rscS* and do not require the gene for squid colonization or competitive fitness. These same strains have a frameshift in *sypE*, and we provide evidence that this Group A *sypE* allele leads to an upregulation in biofilm activity. This work thus describes the central importance of Syp biofilm in colonization of diverse isolates, and demonstrates that significant evolutionary transitions correspond to regulatory changes in the *syp* pathway.

**IMPORTANCE:** Biofilms are surface-associated, matrix-encased bacterial aggregates that exhibit enhanced protection to antimicrobial agents. Previous work has established the importance of biofilm formation by a strain of luminous *Vibrio fischeri* bacteria as the bacteria colonize their host, the Hawaiian bobtail squid. In this study, expansion of this work to many natural isolates revealed that biofilm genes are universally required, yet there has been a shuffling of the regulators of those genes. This work provides evidence that even when bacterial behaviors are conserved, dynamic regulation of those behaviors can underlie evolution of the host colonization phenotype. Furthermore, this work emphasizes the importance of investigating natural diversity as we seek to understand molecular mechanisms in bacteria.

## INTRODUCTION

A fundamental question in studying host-associated bacterial communities is understanding how specific microbial taxa assemble reproducibly in their host. Key insights into these processes were first obtained by studying plant-associated microbes, and the discovery and characterization of Nod factors in Rhizobia was valuable to understand how partner choice between microbe and host could be mediated at the molecular level (1, 2). There are complex communities in humans and other vertebrate animals, yet metagenomic and imaging analyses of these communities have revealed striking reproducibility in the taxa present and in the spatial arrangement of those taxa (3–5). Invertebrate animal microbiomes provide appealing systems in which to study microbiome assembly in an animal host: the number of taxa are relatively small, and examination and manipulation of these organisms have yielded abundant information about processes underlying host colonization (6). For this work we focused on the binary symbiosis between *Vibrio fischeri* and bobtail squids, including the Hawaiian bobtail squid, *Euprymna scolopes*. Bobtail squid have an organ for the symbiont termed the light organ, and passage of specific molecules between the newly-hatched host and the symbiont leads to light organ colonization specifically by planktonic *V. fischeri* and not by other bacteria (7–9). The colonization process involves initiation, accommodation, and persistence steps, resulting in light organ crypt colonization by *V. fischeri*. Upon colonization of the squid light organ, bacteria accumulate to high density and produce light. The bacterial light is modulated by the host to camouflage the moonlight shadow produced by the nighttime foraging squid in a cloaking process termed counter-illumination (10, 11). A diel rhythm leads to a daily clearing of 90-95% of the bacteria from the crypts and regrowth of the remaining cells (12). However, the initial colonization process, including biofilm-based aggregation on the host ciliated appendages, occurs only in newly-hatched squid. This work examines regulation of biofilm formation in diverse squid-colonizing *V. fischeri* strains.

In the well-studied *V. fischeri* strain ES114, biofilm formation is required to gain entry into the squid host. RscS is a hybrid histidine kinase that regulates *V. fischeri* biofilm formation through a phosphorelay involving the hybrid histidine kinase SypF and the response regulator and σ^54^-dependent activator SypG (13–15). This pathway regulates transcription of the symbiosis polysaccharide (Syp) locus, which encodes regulatory proteins (SypA, SypE, SypF, and SypG), glycosyltransferases, factors involved in polysaccharide export, and other biofilm-associated factors (14, 16). The products of the ES114 *syp* locus direct synthesis and export of a biofilm exopolysaccharide that is critical for colonization. Additional pathways have been identified to influence biofilm regulation in ES114, including the SypE-SypA pathway and inhibition of biofilm formation by BinK and HahK (17–21).

*V. fischeri* biofilm regulation is connected to host colonization specificity. In the Pacific Ocean, the presence of *rscS* DNA is strongly correlated to the ability to colonize squid (22). As one example, while the fish symbiont MJ11 encodes a complete *syp* locus, it lacks RscS and does not robustly colonize squid. Heterologous expression of ES114 RscS in MJ11 activates the biofilm pathway and is sufficient to enable squid colonization (22). Similarly, addition of ES114 RscS to *mjapo*.8.1--a fish symbiont that encodes a divergent RscS that is not functional for squid colonization--allows the strain to colonize squid (22). RscS has also been shown to be necessary for squid colonization in certain strains. In addition to ES114, interruption of *rscS* in *V. fischeri* strains KB1A97 and MJ12 renders them unable to colonize squid. Previous phylogenetic analysis revealed that ancestral *V. fischeri* do not encode *rscS*, and that it was acquired once during the organism’s evolution, likely allowing for an expansion in host range. From this analysis, it was concluded that strains with *rscS* can colonize squid, with the only exception being the fish symbionts that harbor the divergent RscS, including *mjapo*.8.1 (22).

There are similar *Vibrio*-squid associations worldwide, yet only *V. fischeri* and the closely-related *Vibrio logei* have been isolated from light organs (23–26). Our 2009 study revealed that although most symbionts have *rscS* DNA, there are Mediterranean *V. fischeri* (e.g., SR5) that do not have *rscS* yet can colonize squid (22, 24, 27). This unexpected finding prompted the current work to examine whether strains such as SR5 colonize with the known biofilm pathway or with a novel pathway. Here, we show that all *V. fischeri* strains tested require the *syp* locus to colonize a squid host, and we identify two groups of isolates that colonize with novel regulation. Given the exquisite specificity by which *V. fischeri* bacteria colonize squid hosts, this work reinforces the importance of biofilm formation and reveals different regulatory modes across the evolutionary tree.

## RESULTS

### Most *V. fischeri* strains synthesize biofilm in response to RscS overexpression

Biofilm formation is required for squid colonization, and overexpression of the biofilm regulator RscS in strain ES114 stimulates a colony biofilm on agar plates (15). Our previous work demonstrated that *V. fischeri* strain MJ11 synthesizes a colony biofilm under similar inducing conditions, which is notable because MJ11 does not encode RscS in its chromosome (22). While the ancestral strain MJ11 did not encode RscS, it had what seemed to be an intact *syp* locus, and overexpression of the heterologous RscS from ES114 was sufficient to enable robust squid colonization (22). We examined a phylogenetic tree of *V. fischeri* isolates (Fig. 1), and in this study we expand our analysis of RscS-Syp biofilm regulation in a wider group of *V. fischeri* strains.

**Figure 1.**
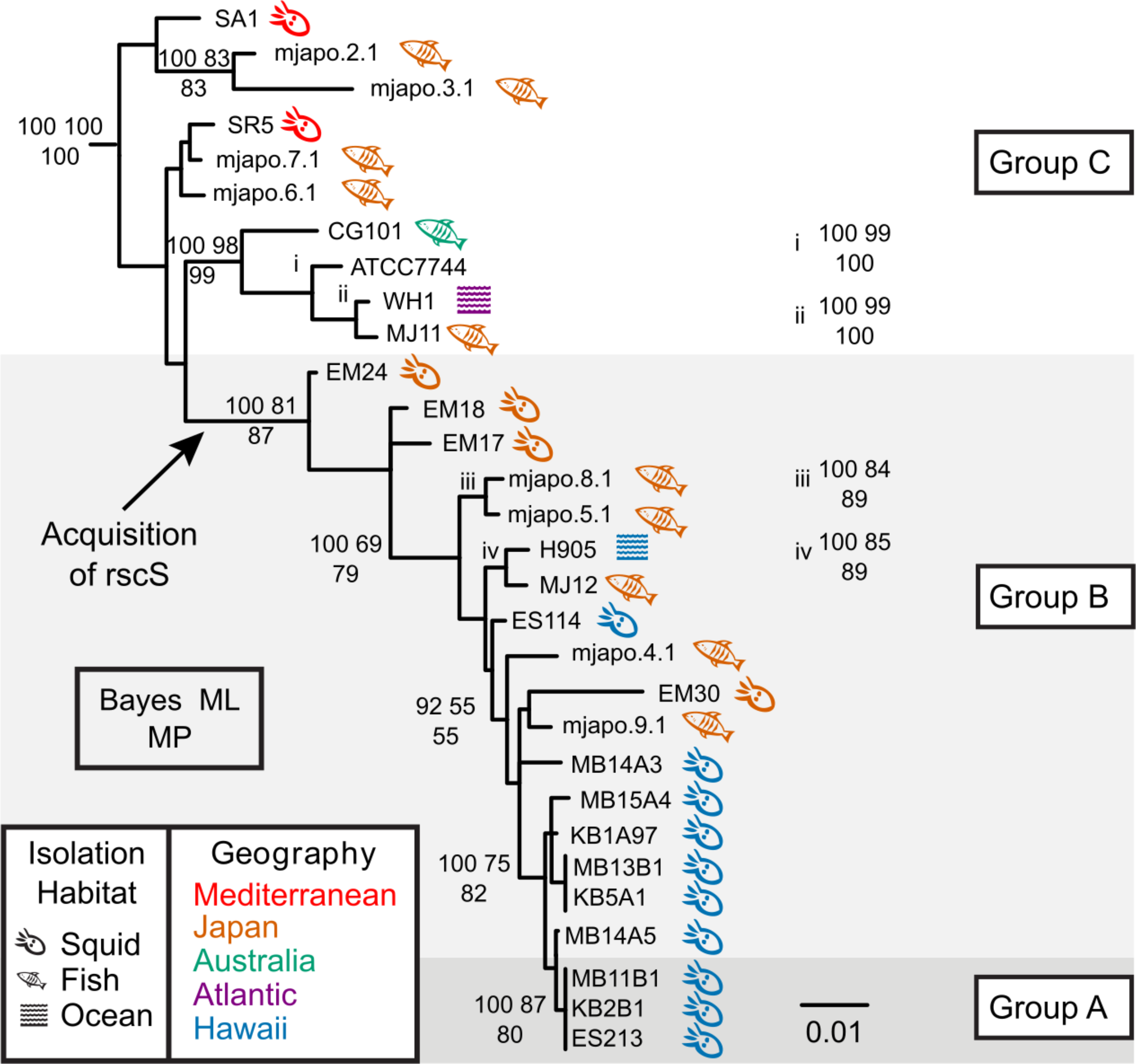
*Vibrio fischeri* phylogeny, highlighting the source of each strain. Bayesian phylogram (50% majority-rule consensus) inferred with a SYM+I+Г model of evolution for the concatenated gene fragments *recA*, *mdh*, and *katA*. In this reconstruction, the root connected to a clade containing the four non-*V. fischeri* outgroup taxa. Statistical support is represented at nodes by the following three numbers: upper left, Bayesian posterior probability (of approximately 37,500 non-discarded samples) multiplied by 100; upper right, percentage of 1000 bootstrap Maximum Likelihood pseudo-replicates; bottom middle center, percentage of 1000 bootstrap Maximum Parsimony pseudo-replicates. Statistical support values are listed only at nodes where more than 2 methods generated support values ≥ 50%. Strains sharing identical sequences for a given locus fragment are listed next to a vertical bar at a leaf; because of a lack of space, some support values have been listed either immediately to the right of their associated nodes and are marked with italicized lower-case Roman numerals in the phylogram. The isolation habitat and geography of each strain are indicated by symbol and color, respectively. The black bar represents 0.01 substitutions/site.

Initially, we asked whether responsiveness to RscS overexpression would yield a similar colony biofilm in this diverse group of strains. We took the same approach as our previous study and introduced plasmid pKG11, which overexpressed ES114 RscS, into strains across the evolutionary tree (22, 28). We observed that almost all strains tested, including those that lack *rscS*, were responsive to overexpression of ES114 RscS (Fig. 2). The morphology of the colony biofilms differed across isolates; but in most cases colony biofilm was evident at 24 h and prominent at 48 h. All of the strains exhibited some wrinkled colony morphology at 48 h with the exception of CG101, which was isolated from the pineapplefish *Cleidopus gloriamaris (25)*. These results demonstrated that most *V. fischeri* strains can produce biofilm in response to RscS overexpression, and this includes strains that presumably have not encountered *rscS* in their evolutionary history.

**Figure 2.**
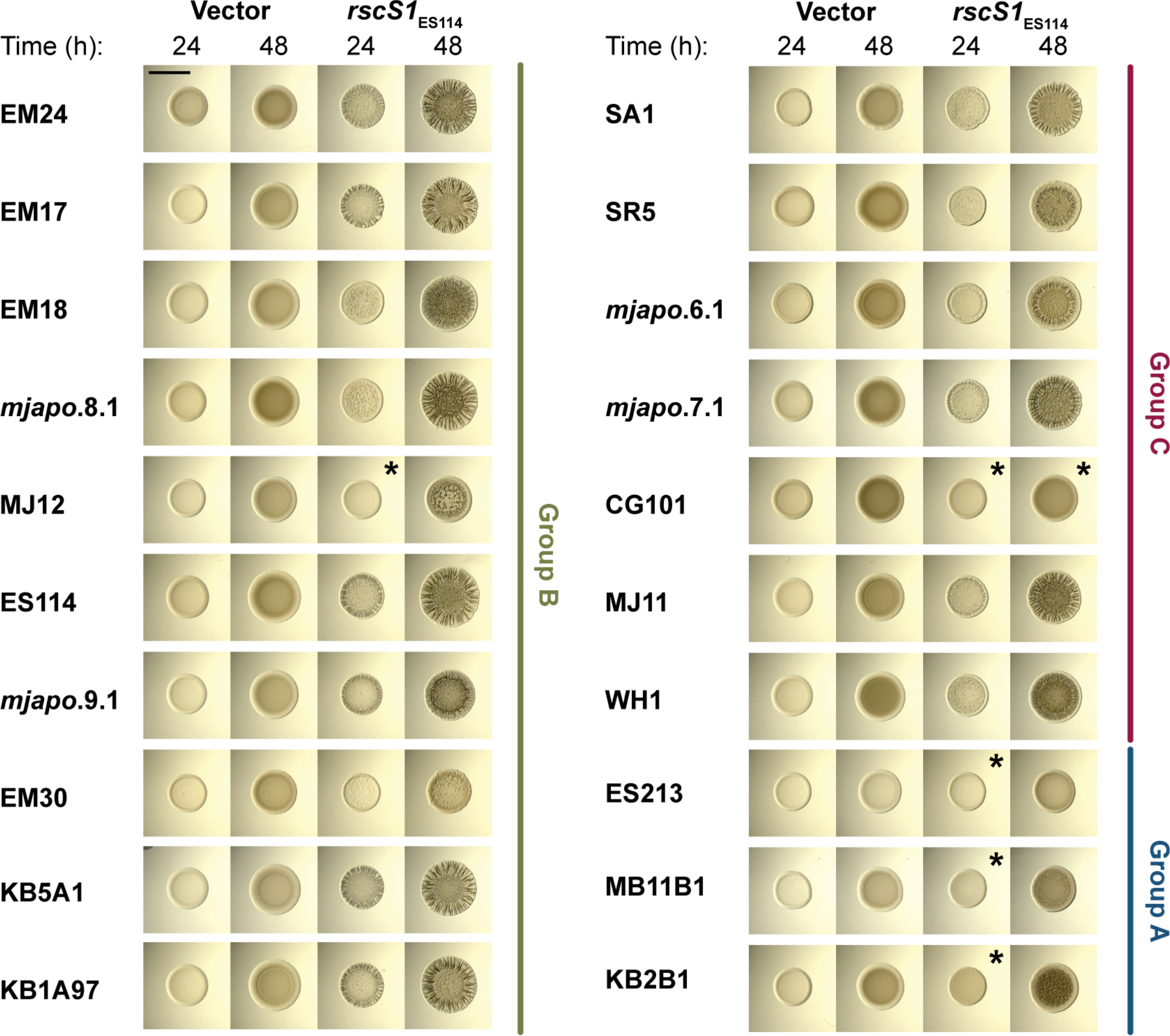
Most *V. fischeri* strains tested form colony biofilm in response to RscS overexpression. Spot assays of the indicated *V. fischeri* strains carrying pKV69 (vector) or pKG11 (*rscS1*; overexpressing ES114 *rscS*) after 24 and 48 h. Strains are MJM1268, MJM1269, MJM1246, MJM1247, MJM1266, MJM1267, MJM1219, MJM1221, MJM1238, MJM1239, MJM1104, MJM1106, MJM1276, MJM1277, MJM1270, MJM1271, MJM1258, MJM1259, MJM1254, MJM1255, MJM1242, MJM1243, MJM1240, MJM1241, MJM1272, MJM1273, MJM1274, MJM1275, MJM1278, MJM1279, MJM1109, MJM1111, MJM1280, MJM1281, MJM1260, MJM1261, MJM1244, MJM1245, MJM1256, and MJM1257. Different phenotypes were observed in the isolates examined; in most cases we observed wrinkled colonies, but in some cases we observed only a subtle pocked pattern (EM30), and in other cases we did not observe any change in colony morphology compared to the vector control (noted by *). The black bar is 5 mm in length.

One unexpected observation was that there was a subset of *rscS*-encoding strains that were reproducibly delayed in their colony biofilm, and had only a mild wrinkled colony phenotype at 48 h (strains MB11B1, ES213, KB2B1; Fig. 2). We considered whether this was due to differential growth of the strains, but resuspension of spots and dilution plating to determine CFU/spot demonstrated no significant growth difference between these strains and ES114 under these conditions. The strains are closely-related (Fig. 1) and a previous study had noted that this group shared a number of phenotypic characteristics, e.g. reduced motility in soft agar (29). Those authors termed this tight clade as “Group A” *V. fischeri* (30). Our results in Figure 2 argue that Group A strains do not respond to RscS in the same manner as other *V. fischeri* strains, which prompted us to investigate the evolution of the RscS-Syp signaling pathway. We have maintained the Group A nomenclature here, and furthermore we introduce the nomenclature of Group B (a paraphyletic group of strains that contain *rscS*; this group includes the common ancestor of all *rscS*-containing strains) and Group C (a paraphyletic group of strains that contains the common ancestor of all *V. fischeri* - these strains do not contain *rscS*), as shown in Figure 1.

### Ancestral Group C squid isolates colonize *E. scolopes* independent of RscS and dependent on Syp

Group C strains generally cannot colonize squid, yet there are Mediterranean squid isolates that appear in this group (Fig. 1; (22)). The best-studied of these strains, SR5, was isolated from *Sepiola robusta*, is highly luminous, and colonizes the Hawaiian bobtail squid *E. scolopes* (24). Nonetheless, this strain lacks *rscS* (27). We first asked whether the strain can colonize in our laboratory conditions, and we confirmed that it colonizes robustly, consistent with the result result previously published by Fidopiastis et al. (24) (Fig. 3). Next, we asked whether it uses the Syp biofilm to colonize. To address this question, we deleted the 18 kb *syp* locus (i.e., *sypA* through *sypR*) in strains SR5 and ES114. Deletion of *rscS* or the *syp* locus in ES114 led to a substantial defect in colonization, consistent with a known role for these factors (Fig. 3). Similarly, deletion of the *syp* locus in SR5, a strain that does not encode *rscS*, led to a dramatic reduction in colonization (Fig. 3). Therefore, even though strain SR5 does not encode *rscS*, it can colonize squid, and it requires the *syp* locus to colonize normally.

**Figure 3.**
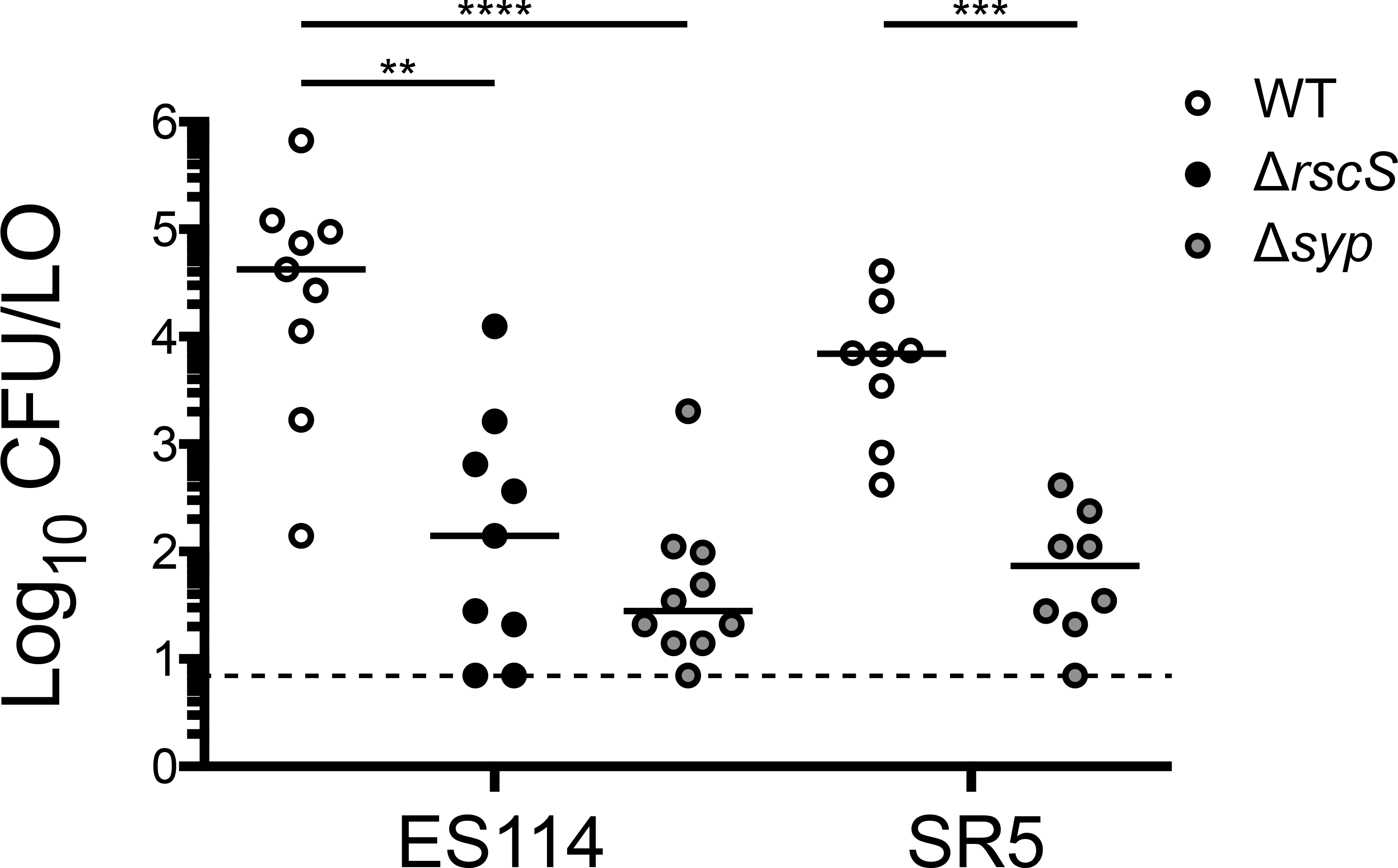
Squid colonization in Group C strain SR5, which does not encode RscS, is dependent on the *syp* polysaccharide locus. Single-strain colonization experiments were conducted and circles represent individual animals. The limit of detection for this assay, represented by the dashed line, is 7 CFU/LO, and the horizontal bars represent the median of each set. Hatchling squid were inoculated with 1.5-3.2 × 10^3^ CFU/ml bacteria, washed at 3 h and 24 h, and assayed at 48 h. Each dot represents an individual squid. Strains are: MJM1100, MJM3010, MJM3062, MJM1125, and MJM3501. Statistical comparisons by the Mann-Whitney test, ** p<0.01, *** p<0.001, **** p<0.0001.

### RscS is dispensable for colonization in Group A strains

We noted in the wrinkled colony biofilm assays shown in Figure 2 that Group A strains exhibited a more modest response to overexpression of RscS. Sequencing of the native *rscS* gene in these strains revealed a predicted −1 frameshift (ΔA1141) between the PAS domain and the histidine kinase CA domain. Whereas ES114 and other Group B strains have nine adenines at this position, the Group A strains have eight, leading to a frameshift and then truncation at an amber stop codon, raising the possibility that Group A strains have a divergent biofilm signaling pathway (Fig. 4A). Given the importance of RscS in the Group B strains including ES114, we considered the possibility that this apparent frameshift encoded a functional protein, either through ribosomal frameshifting or through the production of two polypeptides that together provided RscS function; there is precedent for both of these concepts in the literature (31, 32). We first introduced a comparable frameshift into a plasmid-borne overexpression allele of ES114 *rscS*, and this allele did not function with the deletion of the single adenine (Fig. 4B). This result suggested to us that the frameshift in the Group A strains may not be functional. Therefore, we proceeded to delete *rscS* in two Group A strains (MB11B1, ES213) and two Group B strains (ES114, MB15A4). The Group B strains required RscS for squid colonization (Fig. 5A). However, the Group A strains exhibited no deficit in the absence of *rscS* (Fig. 5A). We next attempted a more sensitive assay in which a Group A strain was competed against MB15A4. Previous studies have demonstrated that in many cases Group A strains outcompete Group B strains (30, 33). We competed Group A strain MB11B1 against Group B strain MB15A4 and observed a significant competitive advantage for the Group A strain, as was observed previously (30). Deletion of *rscS* in the Group A strain did not affect competitive fitness, demonstrating that MB11B1 can outcompete a Group B strain even if MB11B1 lacks RscS (Fig. 5B).

**Figure 4.**
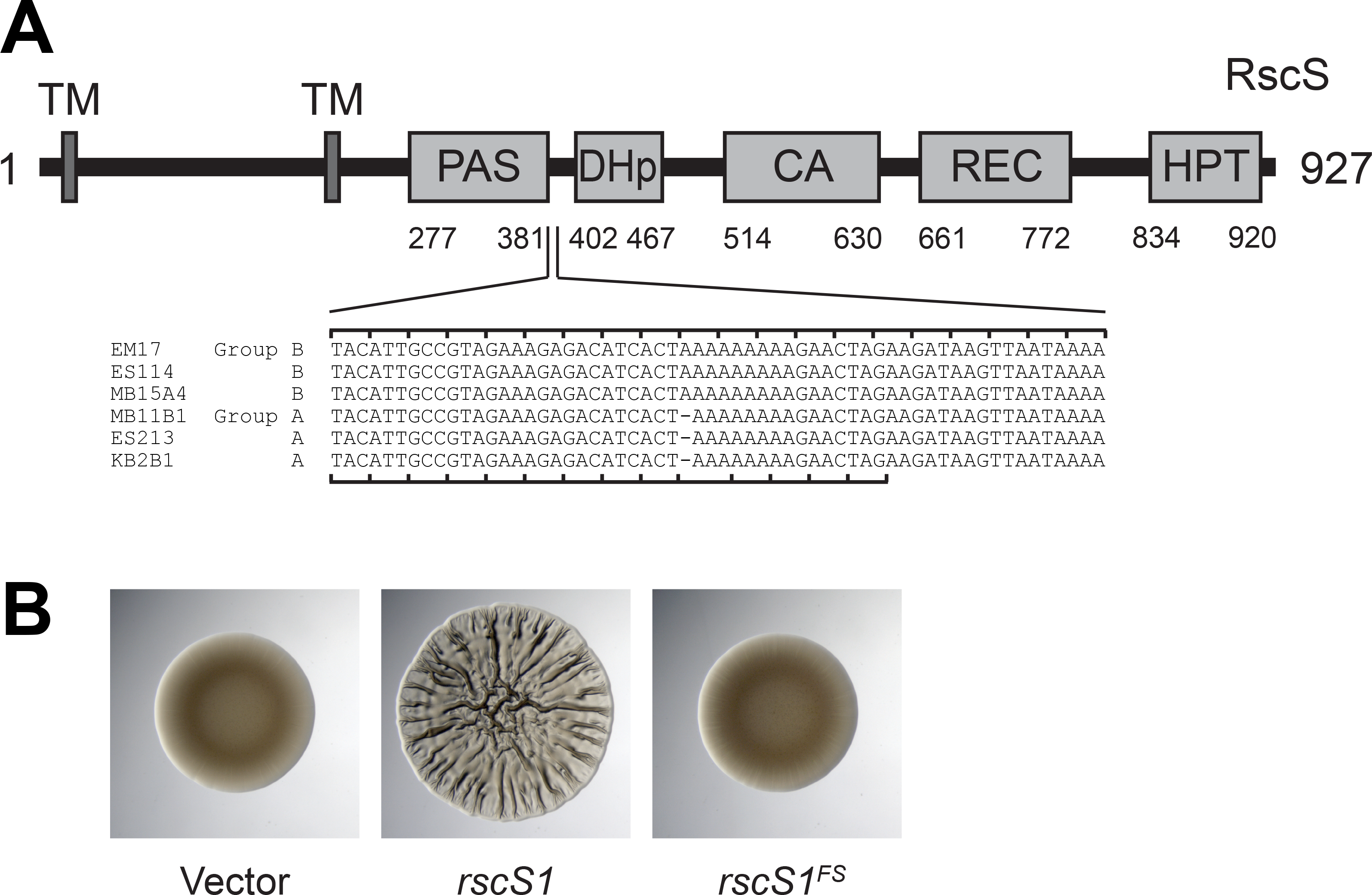
Group A strains have a frameshift in *rscS*. (A) ES114 RscS protein domains. Nucleotides 1114-1173 in ES114 RscS (AF319618) and their homologous sequences in the other Group B and Group A strains are listed. The −1 frameshift is present in the Group A *rscS* alleles. The ES114 reading frame is noted on the top of the alignment and the Group A reading frame on the bottom, which is predicted to end at the amber stop codon. (B) Deletion of nucleotide A1141 in ES114 to mimic this frameshift in pKG11 renders it unable to induce a colony biofilm in a spot assay at 48 h. Strains are MJM1104, MJM1106, and MJM2226.

**Figure 5.**
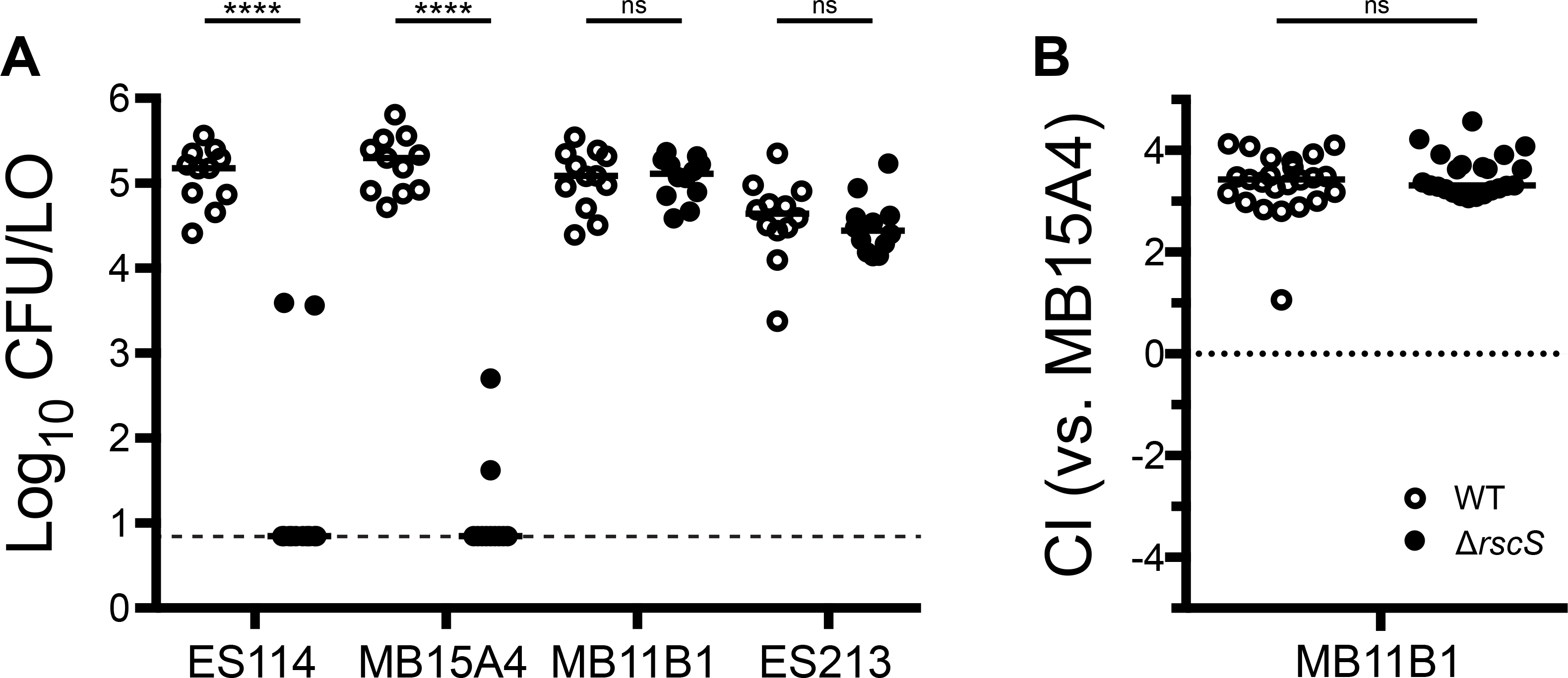
Group A strains MB11B1 and ES213 do not require RscS for squid colonization. Wild-type (WT) and Δ*rscS* derivatives of the indicated strains were assayed in (A) a single-strain colonization assay and (B) competitive colonization against Group B strain MB15A4. Hatchling squid were inoculated at 3.5-14 × 10^3^ CFU/ml bacteria, washed at 3 h and 24 h, and assayed at 48 h. Each dot represents an individual squid. (A) Strains: MJM1100, MJM3010, MJM2114, MJM3042, MJM1130, MJM3046, MJM1117, and MJM3017. The limit of detection is represented by the dashed line, and the horizontal bars represent the median of each set. In both panels, open dots are wild type and filled dots are Δ*rscS*. (B) The competitive index (CI) is defined in the methods and is shown on a Log10 scale. Strains: MJM1130 and MJM3046, each competed against MJM2114. Values greater than 1 indicate more MB11B1. Statistical comparisons by the Mann-Whitney test, ns not significant, **** p<0.0001

### The *syp* locus is broadly required for squid colonization

Given that Group A strains seemed to represent a tight phylogenetic group in which RscS was not required for colonization or competitive fitness, we next asked whether this group requires the Syp biofilm for colonization. We proceeded to delete the entire *syp* locus in two Group A and two Group B strains and to conduct single-strain colonization analysis. In each strain assayed, the *syp* locus was required for full colonization, and we observed a 2-4 log reduction in CFU per animal in the absence of the *syp* genes, pointing to a critical role for Syp biofilm in these strains (Fig. 6). In Group A strains in particular, no colonization was detected in the absence of the *syp* locus. Additionally, we isolated a transposon insertion in SR5 *sypJ*, and this strain did not colonize squid well, arguing that this effect is due to the Syp biofilm and not due to regulation of a distinct phenotype by regulators within the locus.

**Figure 6.**
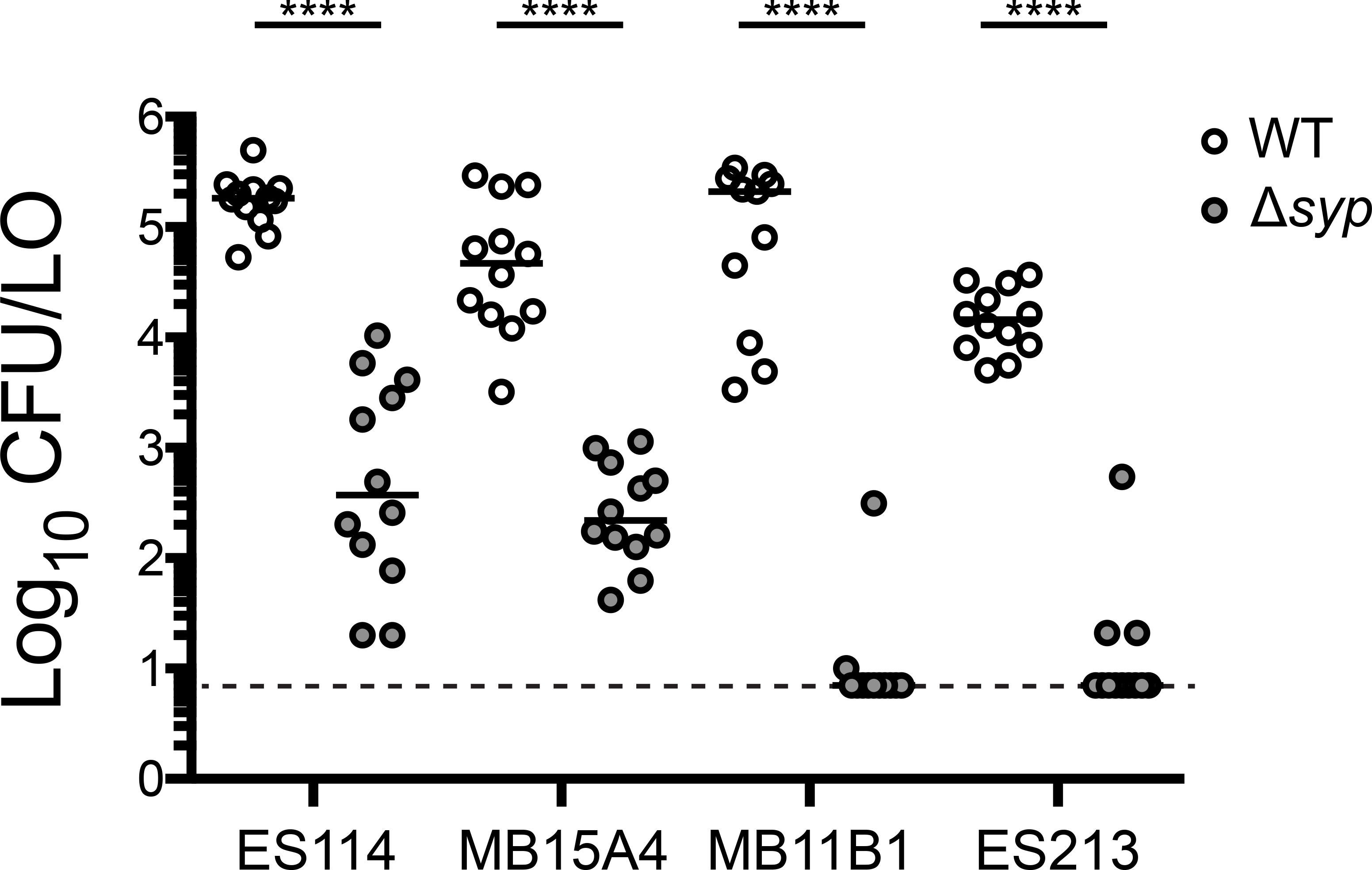
Group B and Group A strains require the *syp* locus for robust squid colonization. Wild type (WT) and Δ*syp* derivatives of the indicated strains were assayed in a single strain colonization assay. Hatchling squid were inoculated with 6.7-32 × 10^2^ CFU/ml bacteria (ES114 and MB15A4 backgrounds) or 5.2-8.9 × 10^2^ CFU/ml bacteria (MB11B1 and ES213 backgrounds), washed at 3 h and 24 h, and assayed at 48 h. Each dot represents an individual squid. The limit of detection is represented by the dashed line and the horizontal bars represent the median of each set. Strains are MJM1100, MJM3062, MJM2114, MJM3071, MJM1130, MJM3065, MJM1117, and MJM3068. Statistical comparisons by the Mann-Whitney test, **** p<0.0001.

### Group A strains encode an alternate allele of SypE

It seemed curious to us that Group A strains do not encode a functional RscS and do not require *rscS* for colonization, yet in many cases Group A strains can outcompete Group B strains (e.g. MB11B1 in Fig. 5B; and Refs. (30, 33)). We reasoned that if the Syp biofilm had a different regulatory architecture in Group A strains--e.g., constitutively activated or activated by a different regulatory protein--then this could explain the Syp regulation independent of RscS. Genome sequencing of SR5 and MB11B1 did not identify a unique histidine kinase that was likely to directly substitute for RscS (27, 33).

Given that the *syp* locus encodes biofilm regulatory proteins, we examined *syp* conservation. We used TBLASTN with the ES114 Syp proteins as queries to determine amino acid conservation in the other *V. fischeri* Group A strain MB11B1, Group C strain SR5, and the *Vibrio vulnificus* type strain ATCC 27562 (34, 35). As shown in Figure 7, ES114 SypE, a response regulator and serine kinase/phosphatase that is a negative regulator of the Syp biofilm (17, 36), exhibited the lowest level of conservation among *syp* locus products. *V. vulnificus* does not encode a SypE ortholog (37), as the syntenic (but not homologous) RbdE encodes a predicted ABC transporter substrate-binding protein. The closest hit for SypE was AOT11_RS12130 (9% identity), compared to 7% identity for the RbdE. Due to the reduced conservation at both the strain and species levels, we analyzed *V. fischeri* MB11B1 SypE in greater detail. Examination of the *sypE* coding sequence revealed an apparent −1 frameshift mutation in which the position 33 (guanine in ES114 and adenine in other Group B and C strains examined) is absent in Group A strains (Fig. 7B). We therefore considered the hypothesis that SypE is nonfunctional in Group A, and that these strains can colonize because they are lacking a functional copy of this negative regulator that is itself regulated by RscS.

**Figure 7.**
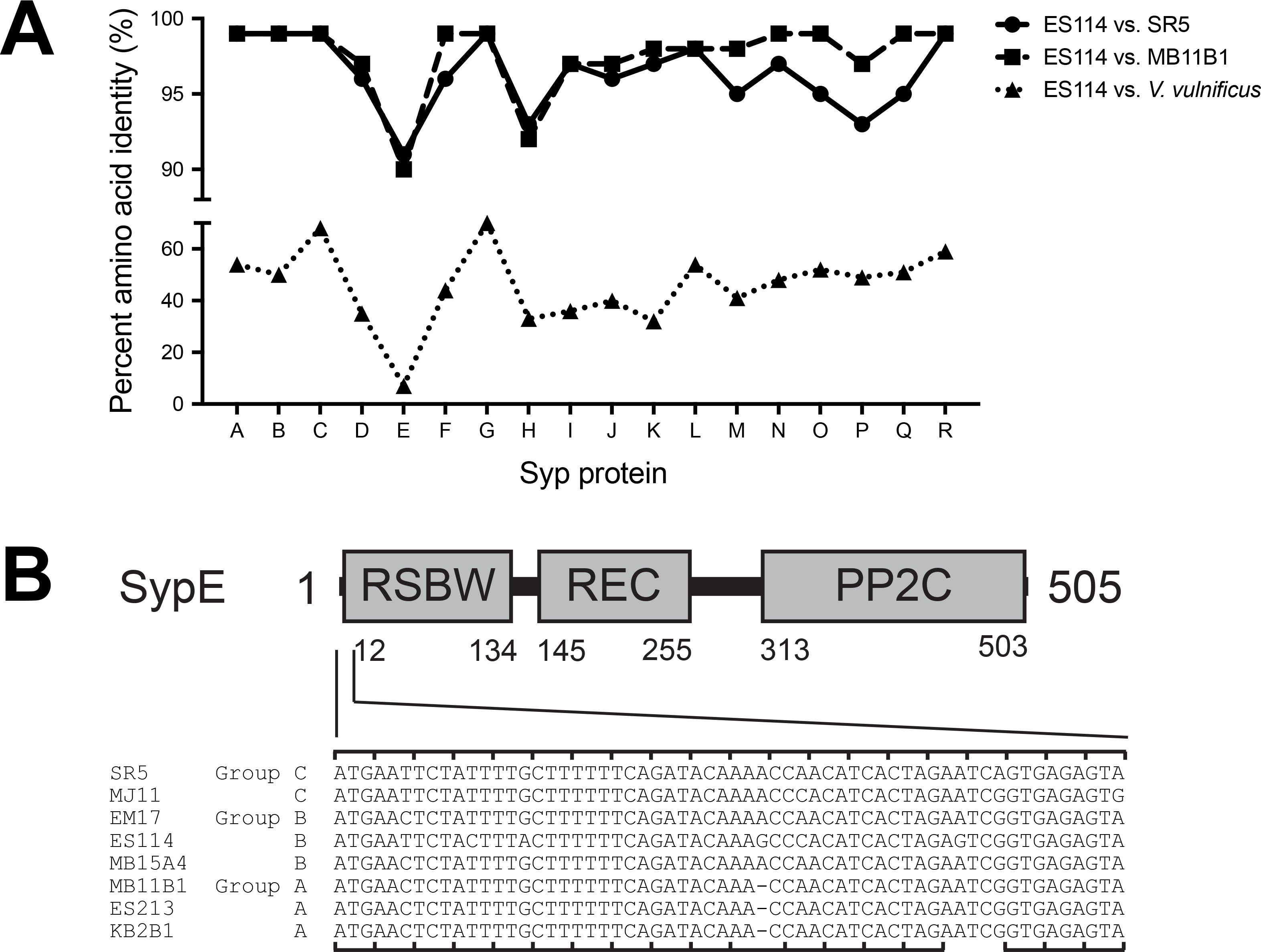
Group A strains have a frameshift in *sypE*. (A) Amino acid identity in the Syp locus. Results show the identity from TBLASTN query using the *V. fischeri* ES114 protein sequences as queries against genes in the homologous loci in *V. fischeri* strains or *V. vulnificus* ATCC 27562. The identity for SypE against *V. vulnificus* is plotted for the syntenous RbdE, although this is not the highest TBLASTN hit, as described in the text. (B) ES114 SypE protein domains. Nucleotides 1-60 in ES114 *sypE* and their homologous sequences in the other Group C, B, and A strains are listed. A −1 frameshift is present in the Group A *sypE* alleles. The ES114 reading frame is noted on the top of the alignment and the Group A reading frame on the bottom, which is predicted to end at the amber stop codon. A possible GTG start codon for the resumption of translation in the ES114 reading frame is present at the position corresponding to the 18th codon in ES114 *sypE*.

To test this hypothesis, we relied on knowledge of the biofilm regulatory pathway from ES114, in which overexpression of SypG produces a wrinkled colony phenotype, but only in strains lacking SypE activity (38, 39). Therefore, we introduced the SypG-overexpressing plasmid pEAH73 into strains as a measure of whether the SypE pathway was intact. In the ES114 strain background, we observed cohesive wrinkled colony formation at 48 h in an ES114 Δ*sypE* strain, but not in the wild-type parent (Fig. 8A). If the *sypE* frameshift observed in MB11B1 led to a loss of function, then introduction of that frameshift into ES114 would lead to a strain that is equivalent to the Δ*sypE* strain. We constructed this strain and upon SypG overexpression we observed wrinkled colony formation. Surprisingly, the biofilm phenotype was observed earlier (i.e., by 24 h) and leads to more defined colony biofilm architecture at 48 h. While the lack of SypE leads to increased and more rapid biofilm formation, in this assay we observed an even greater increase as a result of the frameshift in *sypE* (Fig. 8A).

**Figure 8.**
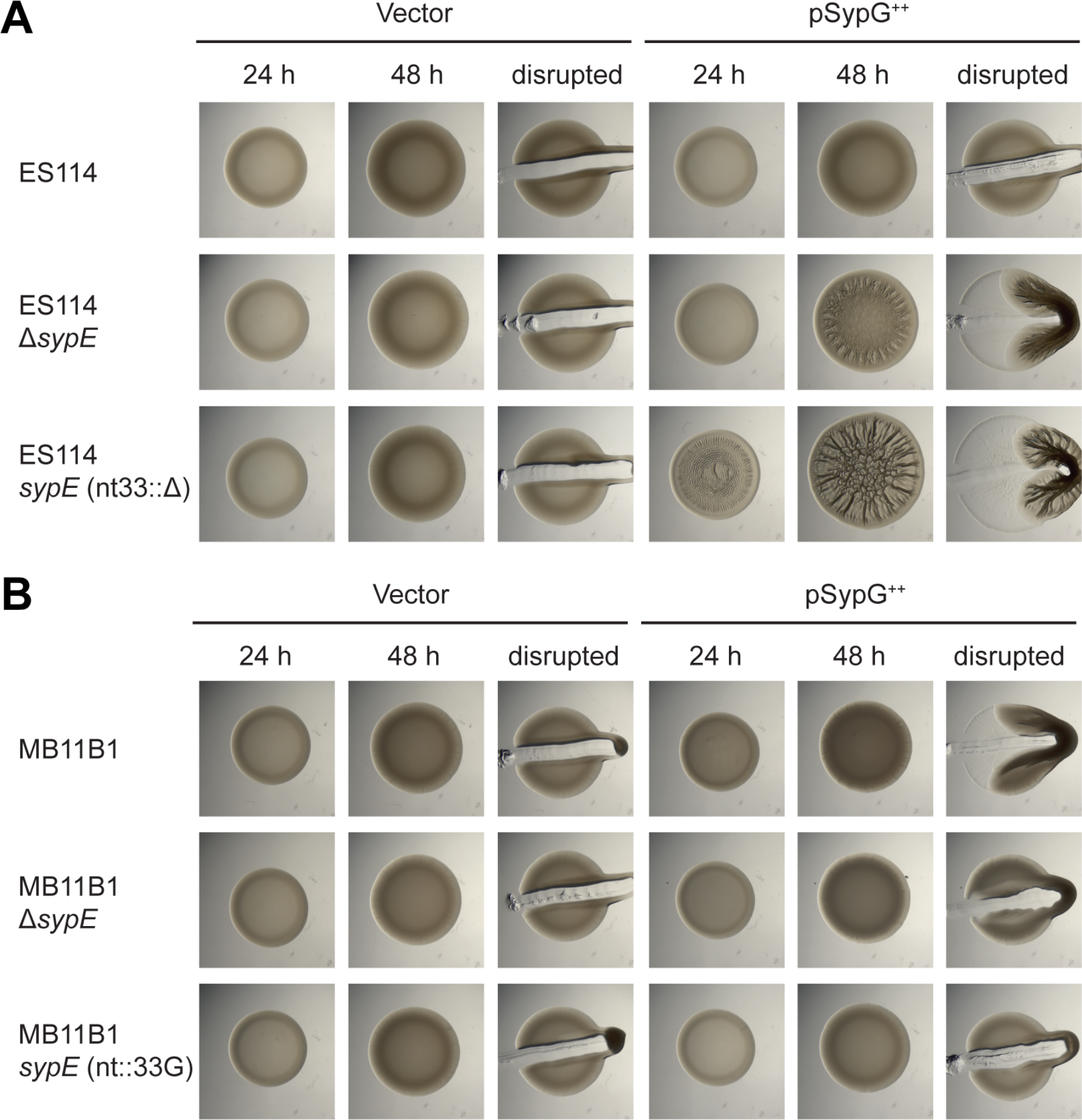
The MB11B1 *sypE* frameshift leads to an enhanced biofilm phenotype upon SypG overexpression. Spot assays of strains carrying the pKV69 vector or pEAH73 SypG overexpression plasmid. (A) ES114 strain background. Strains lacking SypE produce a wrinkled colony phenotype upon SypG overexpression. Deletion of nucleotide 33 in *sypE* to mimic the Group A frameshift led to earlier wrinkling and a more pronounced colony biofilm at 48 h. Strains: MJM1104, MJM3455, MJM3418, MJM3419, MJM3364, and MJM3365. (B) Group A strain MB11B1, which naturally carries a −1 frameshift in *sypE*, exhibits a cohesive phenotype at 48 h with overexpression of SypG. Deletion of *sypE* reduces this phenotype, and repairing the frameshift by addition of a guanosine at nucleotide 33 further reduces the cohesiveness of the spot. Strains: MJM3370, MJM3371, MJM3411, MJM3412, MJM3398, and MJM3399.

We proceeded to conduct a similar assay in the MB11B1 strain background. The colony biofilm phenotypes were muted compared to the ES114 background, but the pattern observed is the same. Strains lacking the additional nucleotide at position 33 (i.e., the native MB11B1 allele) exhibited the strongest cohesion, whereas strains with the nucleotide to mimic ES114 *sypE* (i.e., added back in MB11B1 *sypE*(nt::33G)) were not cohesive (Fig. 8B). These results argue that a novel allele of *sypE* is found in Group A strains and this allele results in more substantial biofilm formation than in a Δ*sypE* strain.

Our finding that the MB11B1 *sypE* allele promotes biofilm formation bolstered the model that this allele contributes to the ability of MB11B1 to colonize squid independent of RscS. To test this model, we introduced the frameshift into ES114 or “corrected” the frameshift in MB11B1. We then conducted single-strain colonization assays, and in each case the *sypE* allele alone was not sufficient to alter the overall colonization behavior of the strain (Fig. 9). Therefore, these data suggest that the frameshift in the MB11B1 *sypE* is not sufficient to explain its ability to colonize independent of RscS, and therefore other regions of SypE and/or other loci in the MB11B1 genome contribute to its ability to colonize independent of RscS.

**Figure 9.**
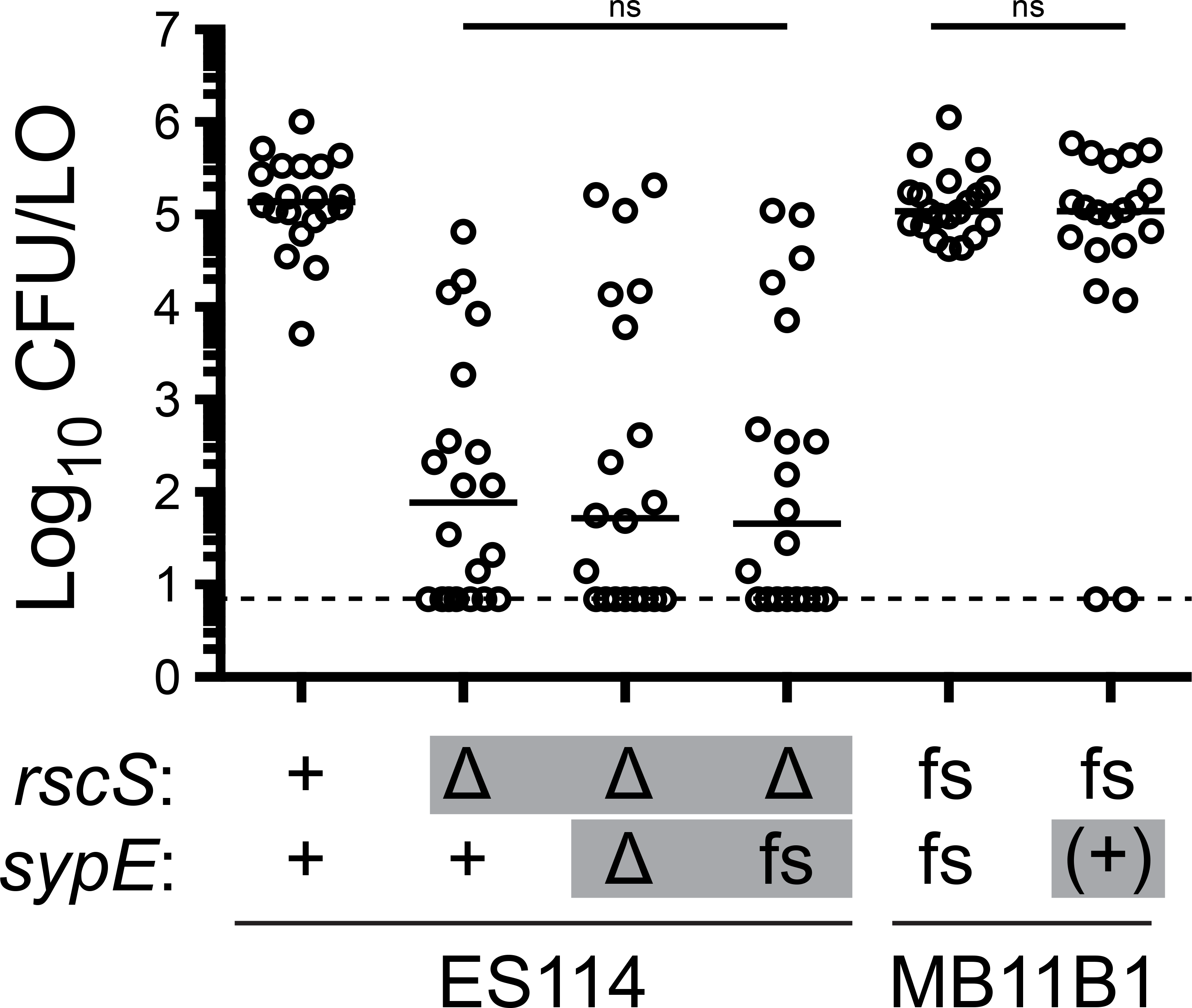
The *sypE*-1 frameshift allele is not sufficient to affect colonization ability. The indicated strains were assayed in a single-strain colonization assay. Gray boxes denote alleles distinct from their wild-type background. Frameshift “fs” refers to alleles--relative to an ES114 reference--that lack *rscS* nucleotide A1141, or that lack *sypE* nucleotide G33. The wild-type MB11B1 strain contains natural frameshifts in these loci, and the ES114 nt33::ΔG allele was constructed. Addition back of the nucleotide in MB11B1 *sypE* is denoted as “(+)”. Hatchling squid were inoculated with 6.8-8.4 × 10^2^ CFU/ml bacteria (MB11B1 background) or 4.0-5.4 × 10^3^ CFU/ml bacteria (ES114 background), washed at 3 h and 24 h, and assayed at 48 h. Each dot represents an individual squid. The limit of detection is represented by the dashed line and the horizontal bars represent the median of each set. Strains are MJM1100, MJM3010, MJM4323, MJM3394, MJM1130, and MJM3397. Statistical comparisons by the Mann-Whitney test, ns not significant

### BinK is active in Group A, B, and C strains

We recently described the histidine kinase, BinK, which negatively regulates *syp* transcription and Syp biofilm formation (18). In ES114, overexpression of BinK impairs the ability of *V. fischeri* to colonize. We therefore reasoned that if BinK could function in Group A strains and acted similarly to repress Syp biofilm, then overexpression of BinK would reduce colonization of these strains. We introduced the pBinK plasmid (i.e., ES114 *binK* (18)) and asked whether multicopy *binK* would affect colonization. In strain MB11B1, BinK overexpression led to a dramatic reduction in colonization (Fig. 10A). Therefore, there is a clear effect for BinK overexpression on the colonization of the Group A strain MB11B1.

**Figure 10.**
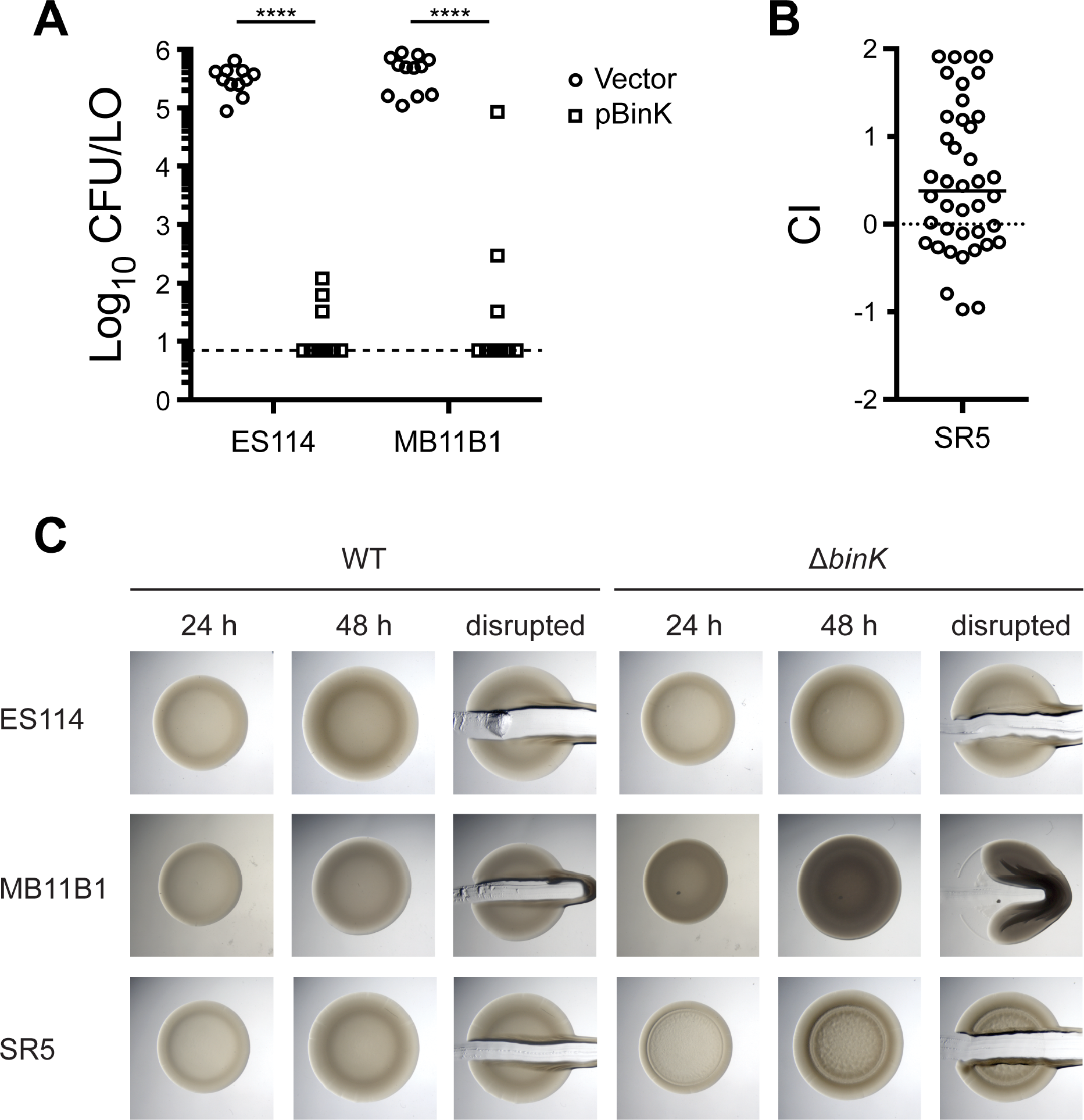
BinK is active in Groups A, B, and C. (A) Overexpression of pBinK inhibits colonization in Group A strain MB11B1. Hatchling squid were inoculated with 3.6-6.8 × 10^3^ CFU/ml bacteria, washed at 3 h and 24 h, and assayed at 48 h. Each dot represents an individual squid. The limit of detection is represented by the dashed line and the horizontal bars represent the median of each set. The vector control is pVSV104. Strains are MJM1782, MJM2386, MJM2997, and MJM2998. (B) Deletion of *binK* confers a colonization defect in Group C strain SR5. Strains are MJM1125 and MJM3571; mean inoculum of 7.2 × 10^3^ CFU/ml; median competitive index (CI) was 0.38 (i.e., 2.4-fold advantage for the mutant). (C) Deletion of the native *binK* in MB11B1 yielded opaque and cohesive spots, which are stronger phenotypes than we observe in ES114. Strains are MJM1100, MJM2251, MJM1130, MJM3084, MJM2997, and MJM2998. Statistical comparisons by the Mann-Whitney test, **** p<0.0001.

We attempted to ask the same question in Group C strain SR5, but the pES213-origin plasmids were not retained during squid colonization. Therefore, we instead asked whether deletion of the BinK, a negative regulator of ES114 colonization, has a comparable effect in SR5 (18). We deleted *binK* and observed a 2.4-fold competitive advantage during squid competition (Fig. 10B), arguing that BinK in this Group C strain is active and performs an inhibitory function similar to that in ES114.

We next examined the colony biofilm phenotype for strains lacking BinK. MB11B1 Δ*binK* exhibited a mild colony biofilm phenotype at 48 h, as evidenced by the cohesiveness of the spot when disrupted with a toothpick (Fig. 10C). The colonies also exhibited an opaque phenotype. In a minority of experimental replicates, wrinkled colony morphology was evident at 48 h, but in all samples wrinkled colony morphology was visible at 7 d (data not shown). The SR5 Δ*binK* strain also exhibited slightly elevated biofilm morphology at 48 h, though the cells were not as cohesive as those of MB11B1 Δ*binK* (Fig. 10C). Together, the results in Figure 10 argue that BinK, a factor that has been characterized as a negative regulator of Syp biofilm, plays similar roles in Group A and Group C strains and has a widely-conserved function across the *V. fischeri* evolutionary tree.

## DISCUSSION

This study examines regulation of a beneficial biofilm that is critical to host colonization specificity in *V. fischeri.* The Syp biofilm was discovered thirteen years ago and has been characterized extensively for its role in facilitating squid colonization by *V. fischeri.* This work establishes that the *syp* locus is required broadly across squid symbionts, and it uncovers three groups of *V. fischeri* that use different regulatory programs upstream of the *syp* locus. A simplified phylogenetic tree showing key features of squid symbionts in these three groups is shown in Figure 11.

**Figure 11.**
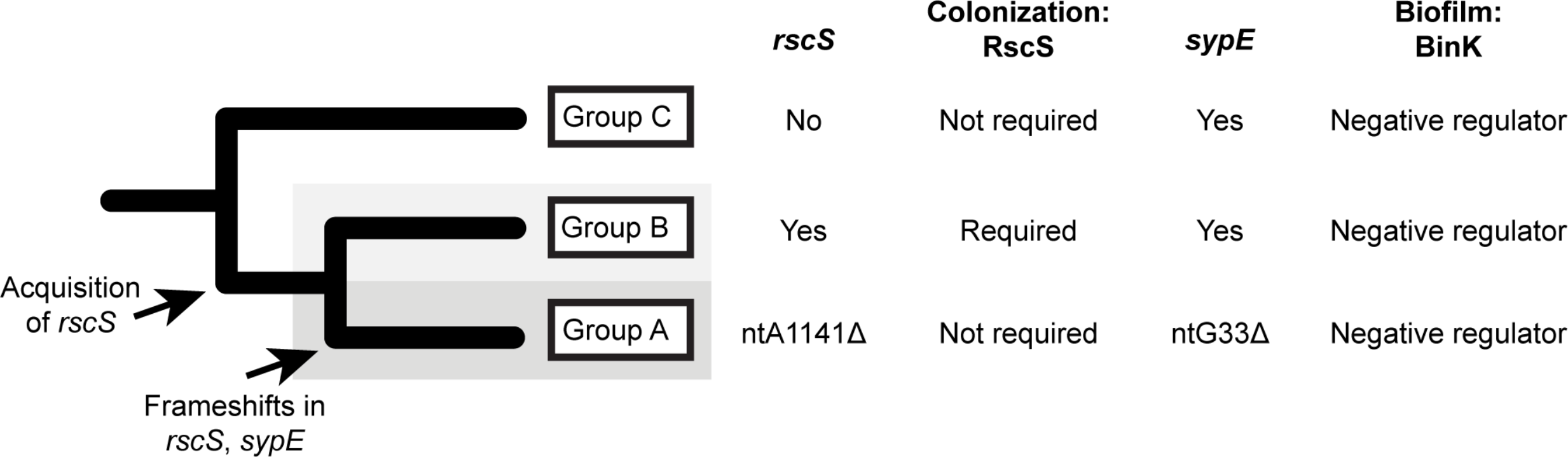
Summary model of distinct modes of biofilm formation in squid-colonizing *V. fischeri*. Phylogenetic tree is simplified from Figure 1, and illustrates key features of squid symbionts in the three groups. Shown are divergent aspects (RscS, SypE) and conserved regulation (BinK). In all groups, the *syp* exopolysaccharide locus is required for squid colonization.

There are three nested evolutionary groups of *V. fischeri* that have been described separately in the literature and here we formalize the nomenclature of Groups C, B, and A. Group A is a monophyletic group, as are Groups AB and ABC (Fig. 1). This work provides evidence that squid symbionts in each group have a distinct biofilm regulatory architecture. Most *V. fischeri* isolates that have been examined from the ancestral Group C cannot colonize squid; however, those that can colonize do so without the canonical biofilm regulator RscS. We show that the known targets of RscS regulation—genes in the *syp* biofilm locus—are nonetheless required for squid colonization by this group. Group B strains include the well-characterized ES114 strain, which requires RscS and the *syp* locus to colonize squid. Group A strains differ phenotypically and behaviorally from the sister Group B strains (30), and we demonstrate that these strains have altered biofilm regulation. Group A strains have a frameshift in *rscS* that renders it nonfunctional, and a 1 bp deletion in *sypE*, and we provide evidence that the *sypE* allele promotes biofilm development in the absence of RscS. Additionally, we note that the *sypE* frameshift is not present in SR5, arguing for distinct modes of biofilm regulation in Groups A, B, and C.

At the same time, this study provides evidence that some aspects of biofilm regulation are conserved in diverse squid symbionts, such as the effects of the strong biofilm negative regulator BinK. Published data indicate that evolved BinK alleles can alter colonization of H905 (Group B) and MJ11 (Group C), and that a deletion of MJ11 *binK* leads to enhanced colonization (20). Our experiments in Figure 10 show a clear effect for BinK in all three phylogenetic groups. We also observed responsiveness to RscS overexpression in all squid symbionts examined (Fig. 2). CG101 was the only *V. fischeri* strain examined that did not exhibit a colony biofilm in response to RscS overexpression. CG101 was isolated from the Australian fish *Cleidopus gloriamaris*; based on these findings, we suspect that the strain does not have an intact *syp* locus or otherwise has divergent biofilm regulation.

It remains a formal possibility that the entire *syp* locus is not required in Group A or Group C, but instead that only one or a subset of genes in the locus are needed. We have constructed Campbell-type (insertion duplication) alleles to interrupt *sypG* in MB11B1 and SR5, and additionally have isolated a transposon insertion in SR5 *sypJ*, and none were able to colonize well. Additionally, aggregation in squid mucus has been observed for the Group A strain MB13B2, and this aggregation is dependent on *sypQ* (40). In our data we note that Group A strains were completely unable to colonize in the absence of the *syp* locus, unlike the tested Group B&C strains that exhibited reduced colonization in their respective mutants (Figs. 3, 6). Therefore, the simplest explanation is that the *syp* locus is required in divergent strains in a manner similar to how it is used in ES114. We think that the ability to completely delete the *syp* locus is a clean way to ask whether the locus is required for specific phenotypes, and our strains are likely to be useful tools in probing Syp protein function in diverse *V. fischeri* isolates.

It is intriguing to speculate as to how the two frameshifts in the Group A strains arose, and why the nonfunctional RscS is tolerated in this group. One possible scenario is that the Group A strains acquired a new regulatory input into the Syp pathway, and that the presence of this new regulator bypassed the requirement for RscS. We note that comparative genomic analysis of Hawaiian D (dominant)-type strains--which largely overlap with Group A--revealed an additional 250 kb of genomic DNA compared to other isolates, yielding a large cache of genes that could play a role in this pathway (33). A related possibility is that *rscS*-independent colonization results from altered regulation of the *syp* locus, either due to changes in regulators (e.g. SypF) or sites that are conserved with Group B. An additional possibility is that the *sypE* frameshift arose, enabling Group A strains to colonize independent of *rscS*. Given that correction of this frameshift in MB11B1 does not significantly affect colonization ability (Fig. 9), this sequence of events seems less likely, and we expect that another regulator in MB11B1 is required for the RscS-independent colonization phenotype. There is evidence that under some conditions LuxU can regulate the *syp* biofilm (41), and as this protein is conserved in *V. fischeri* it may play an important role in Group A or Group C.

Results from two experimental conditions suggest that the Group A strains may have an elevated baseline level of biofilm formation. Our data indicate that in the absence of BinK or upon SypG overexpression, MB11B1 colonies exhibit strong cohesion under conditions in which ES114 does not (Figs. 8, 10). Furthermore, we note that the Group A strain MB11B1, when lacking BinK, also exhibits a darker, or more opaque, colony phenotype (Fig. 10). This phenotype has been observed in some ES114 mutants (16) but not in the corresponding ES114 Δ*binK* strain (Fig. 10). The entire colonization lifecycle likely requires a balance between biofilm formation/cohesion and biofilm dispersal, and these data argue that Group A strains may be more strongly tilted toward the biofilm-producing state. There is evidence that strains lacking BinK exhibit a colonization advantage in the laboratory (18, 20), suggesting that this strategy of more readily forming biofilms may provide a fitness advantage in nature. At the same time, the biofilm negative regulator BinK is conserved among *V. fischeri* strains examined (including MB11B1; Fig. 10), arguing that there is a benefit to reducing biofilm formation under some conditions.

Our study provides hints as to the role of SypE in MB11B1 and other Group A strains. In ES114, the C-terminus is a PP2C serine kinase domain, whereas the N-terminus of SypE is an RsbW serine phosphatase domain. SypE acts to phosphorylate and dephosphorylate SypA Ser-56, with the unphosphorylated SypA being the active form to promote biofilm development (17). The balance between SypE kinase and phosphatase is modulated by a central two-component receiver domain (17). Our data that the MB11B1 *sypE* allele promotes biofilm formation suggest that the protein is tilted toward the phosphatase activity. In MB11B1, the frameshift early in *sypE* suggests that there is a different start codon and therefore a later start codon. An alternate GTG start codon in MB11B1 occurs corresponding to codon 18 in ES114 *sypE* (Fig. 7), and this is likely the earliest start for the MB11B1 polypeptide. We attempted to directly identify the SypE N-terminus by mass spectrometry, yet we could not identify the protein from either strain. Additional study is required to elucidate how MB11B1 SypE acts to promote biofilm formation. *V. fischeri* strains are valuable symbionts in which to probe the molecular basis to host colonization specificity in animals (22, 25, 26). A paradigm has emerged in which biofilm formation through the RscS-Syp pathway is required for squid colonization but not for fish colonization. This study affirms a role of the Syp biofilm, but at the same time points out divergent (RscS-independent) regulation in Group C and Group A isolates. In another well-studied example of symbiotic specificity, Rhizobial Nod factors are key to generating specificity with the plant host, yet strains have been identified that do not use this canonical pathway (42, 43). Future work will elaborate on these RscS-independent pathways to determine how non-canonical squid colonization occurs in diverse natural isolates.

## MATERIALS&METHODS

### Bacterial strains and growth conditions

*V. fischeri* and *E. coli* strains used in this study can be found in Table 1. *E. coli* strains, used for cloning and conjugation, were grown in Luria-Bertani (LB) medium (25 g Difco LB Broth [BD] per liter). *V. fischeri* strains were grown in Luria-Bertani salt (LBS) medium (25 g Difco LB Broth [BD], 10 g NaCl, and 50 ml 1 M Tris buffer pH 7.0, per liter). Growth media were solidified by adding 15 g Bacto agar (BD) per liter. When necessary, antibiotics (Gold Biotechnology) were added at the following concentrations: tetracycline, 5 μg/ml for *V. fischeri*; erythromycin, 5 μg/ml for *V. fischeri*; kanamycin, 50 μg/ml for *E. coli* and 100 μg/ml for *V. fischeri*; and chloramphenicol, 25 μg/ml for *E. coli*, 2.5 −5 μg/ml for Group B *V. fischeri*, and 1 - 2.5 μg/ml for Group A *V. fischeri*. The two MB11B1 / pKV69 strains listed reflect two separate constructions of this strain, though we have not identified any differences between them.

**Table 1.**
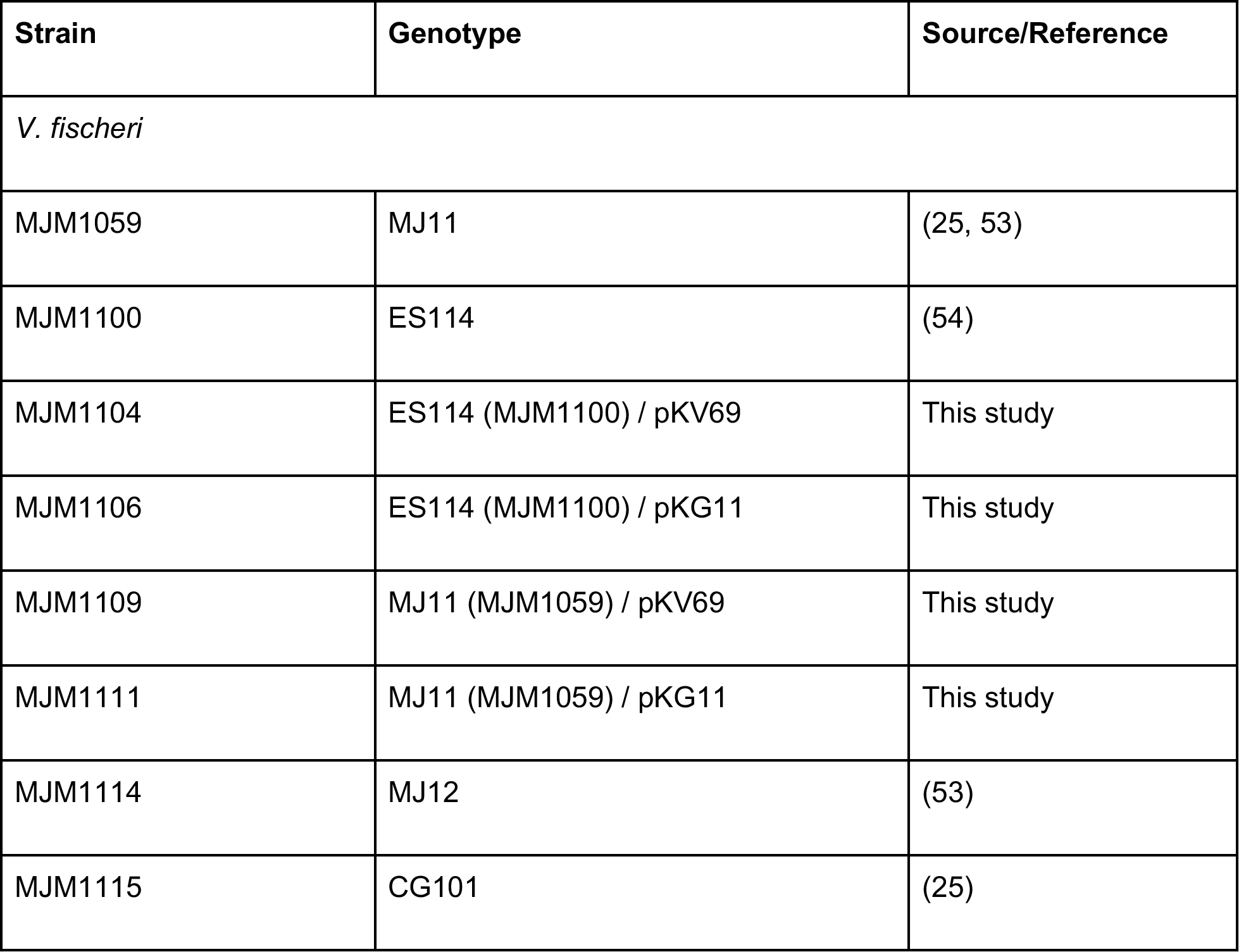
Bacterial strains

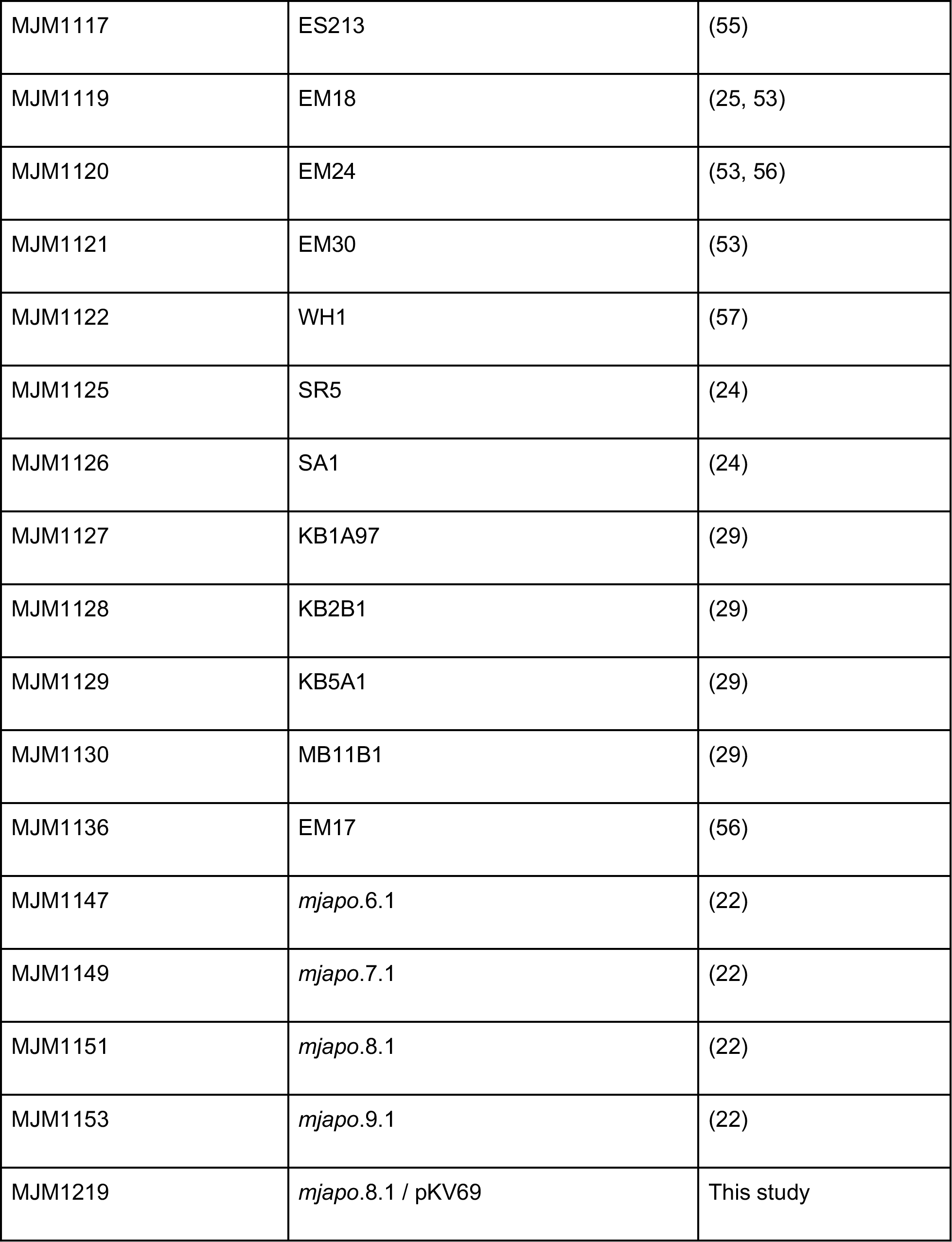

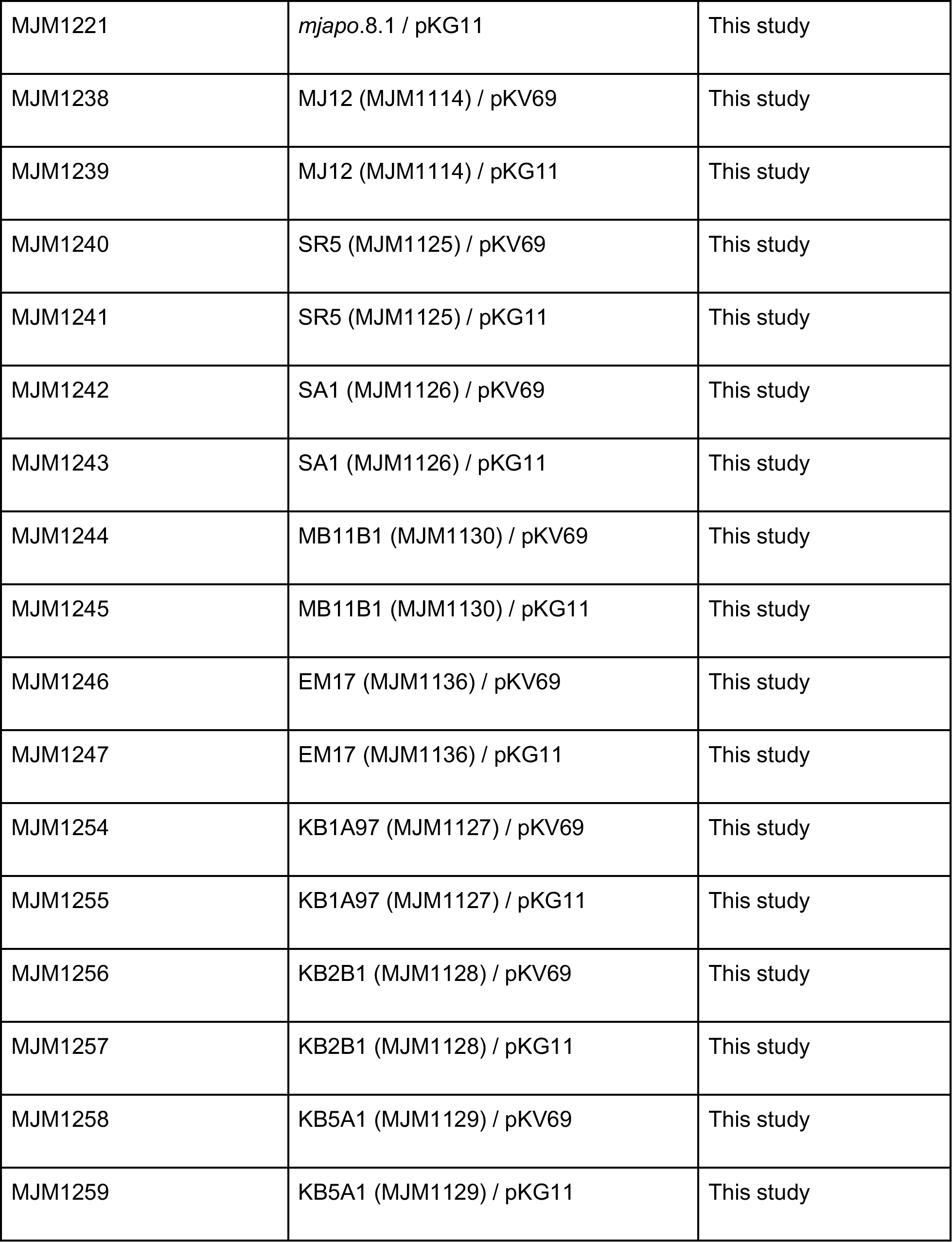

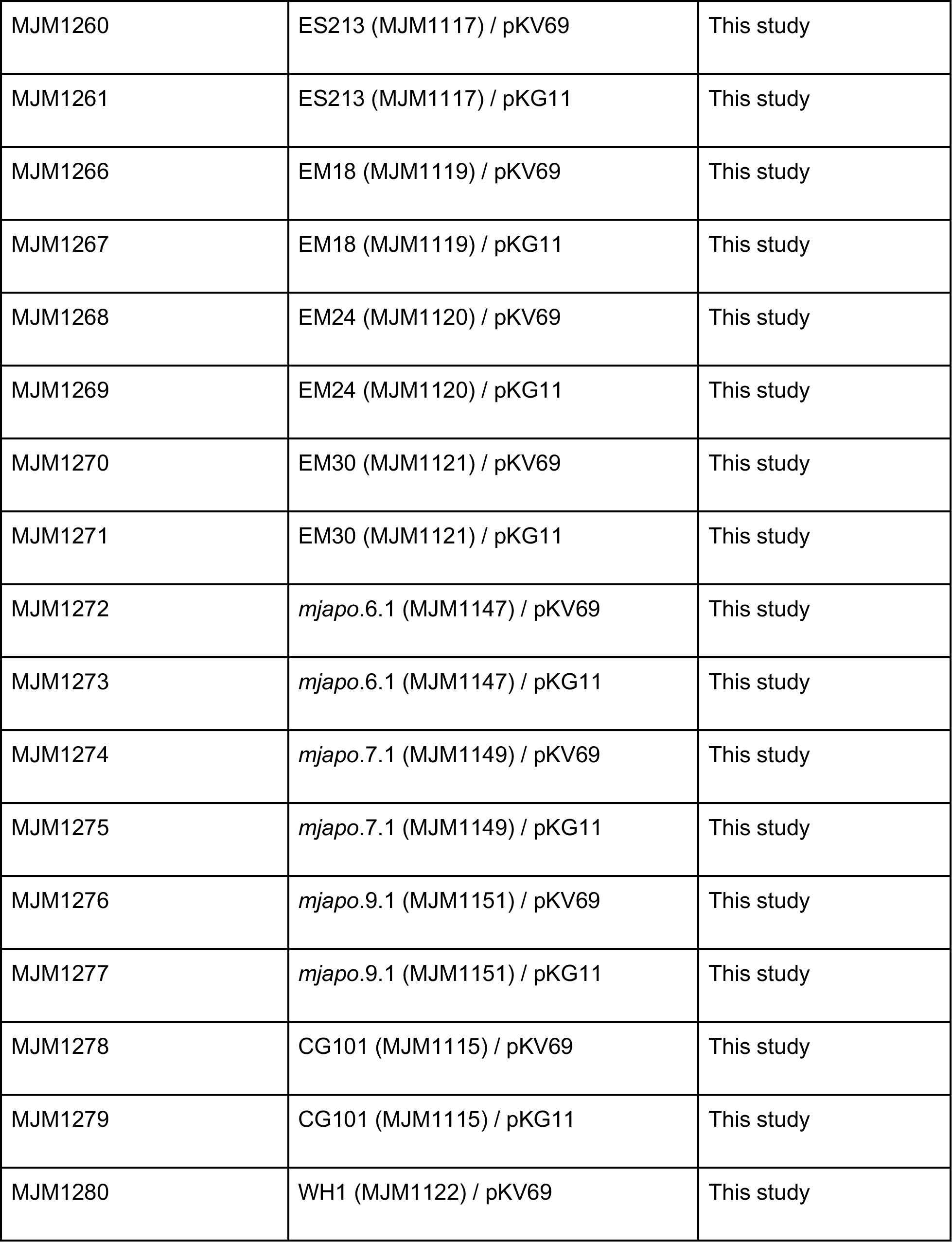

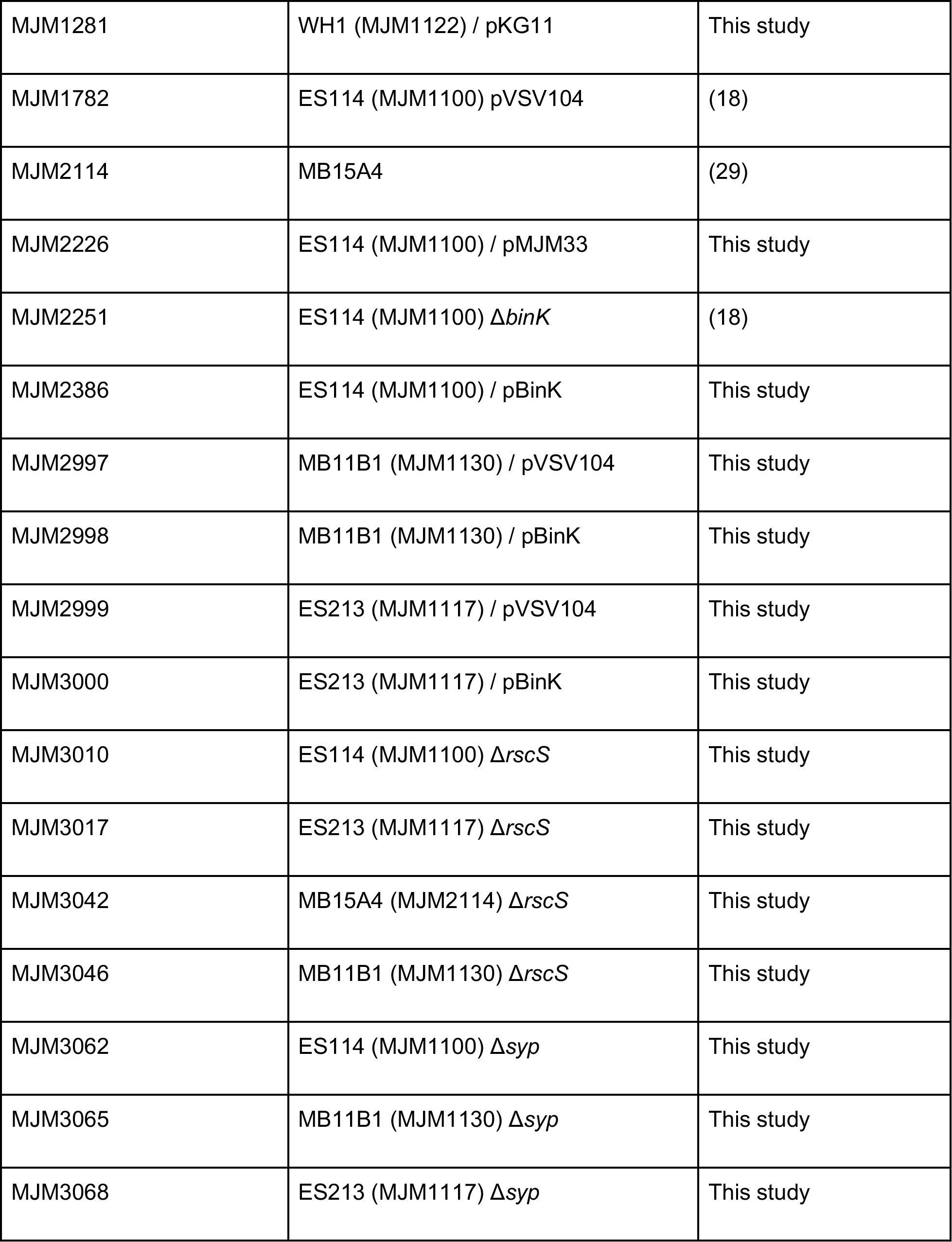

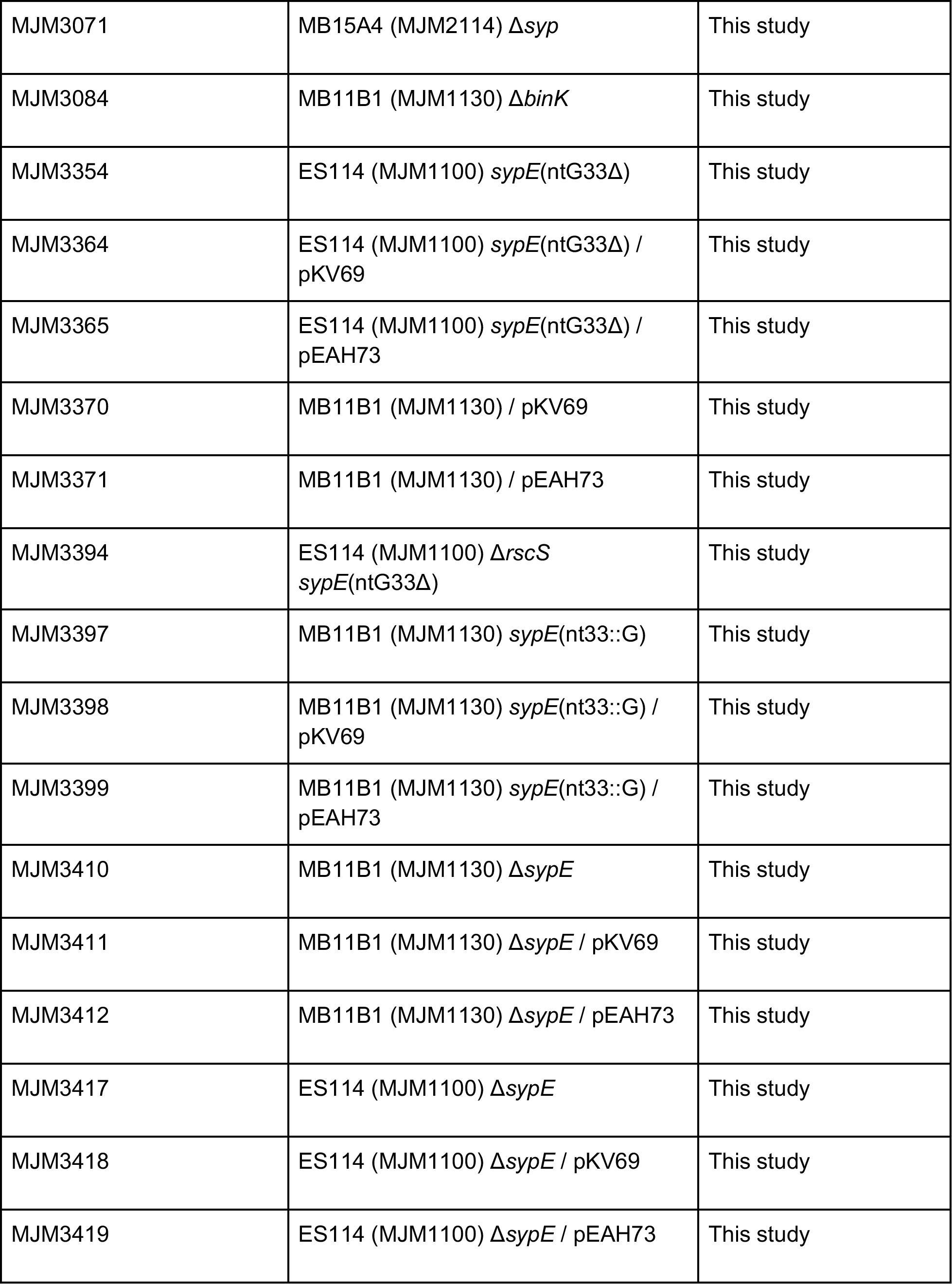

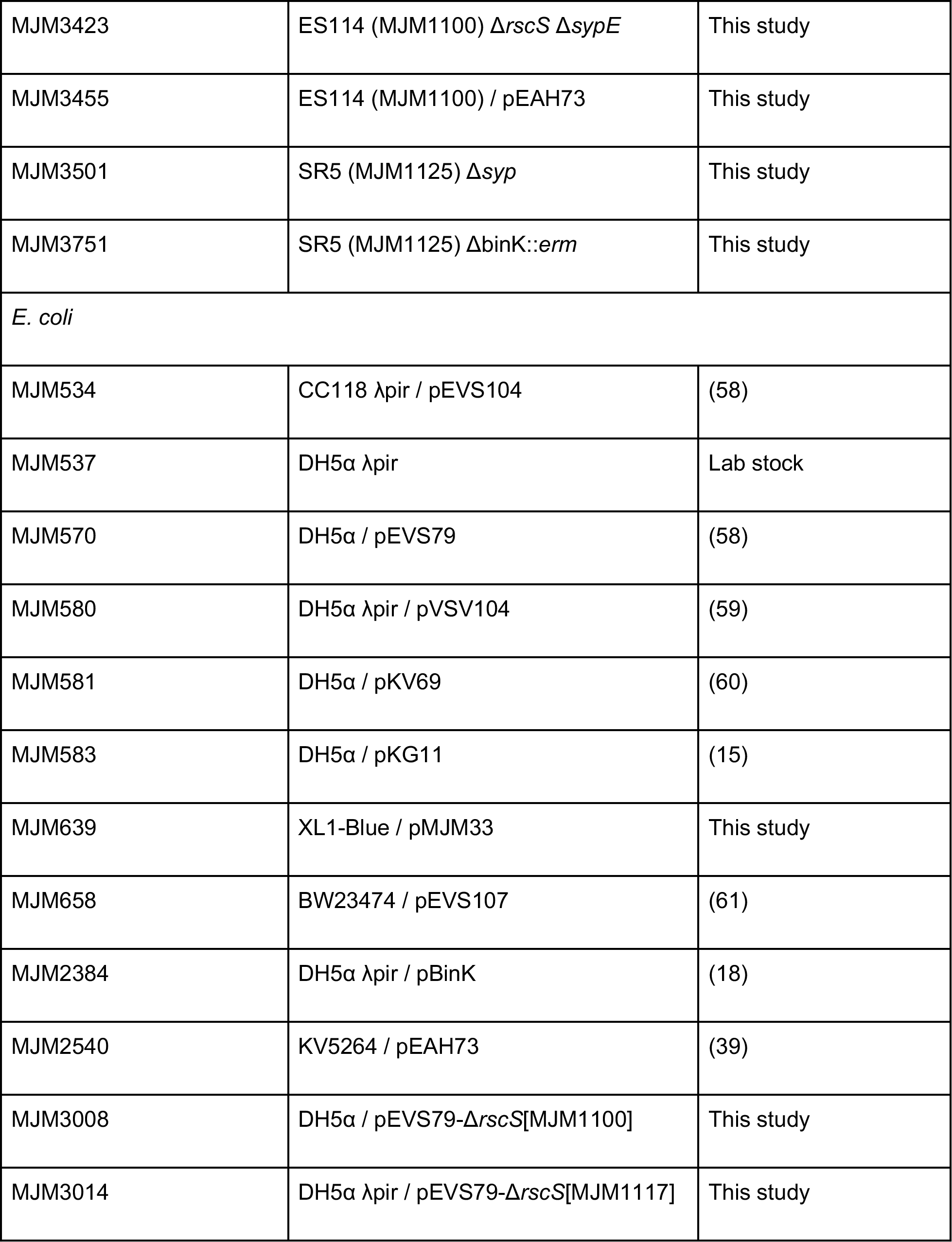

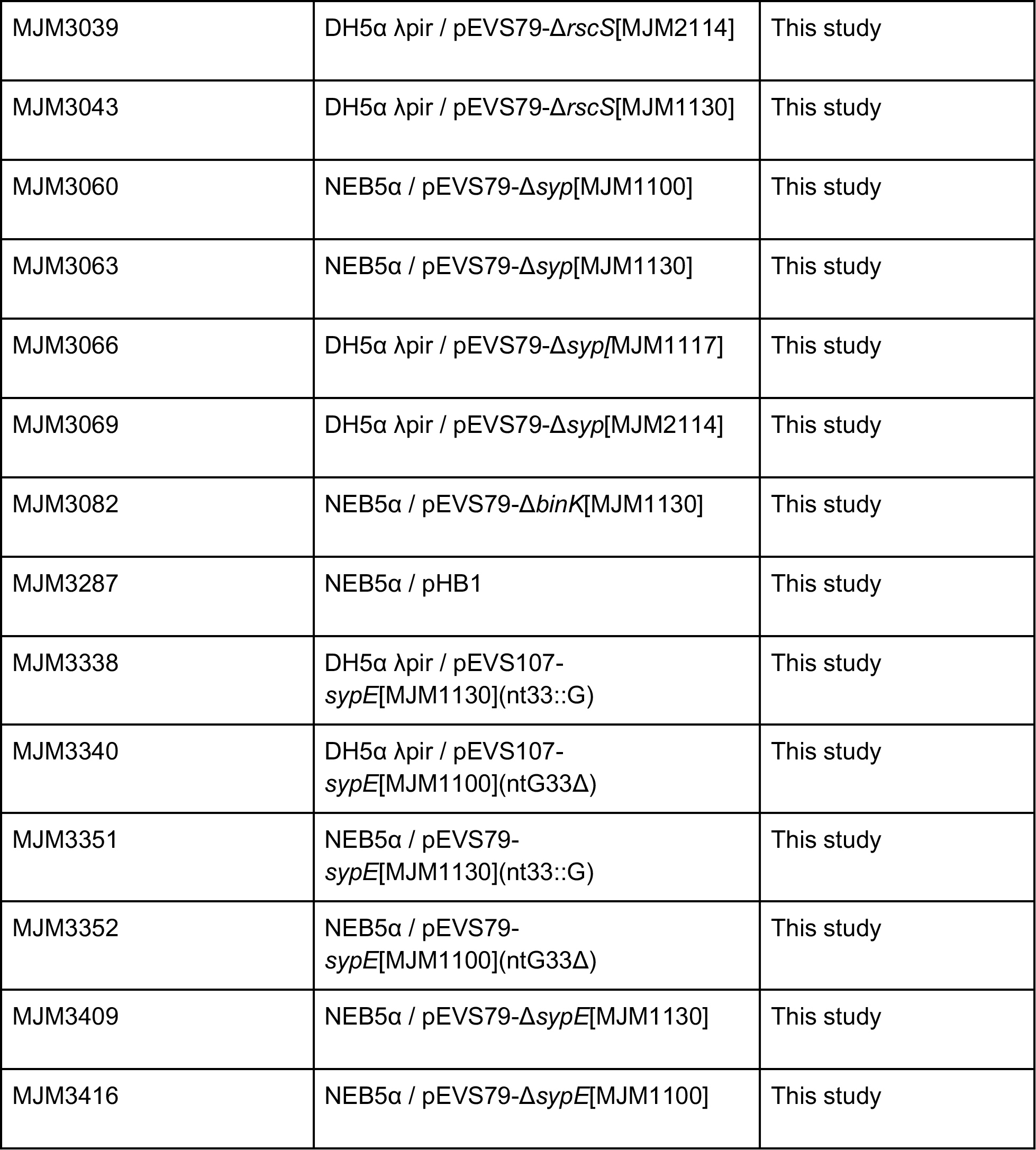

**Table 2.**
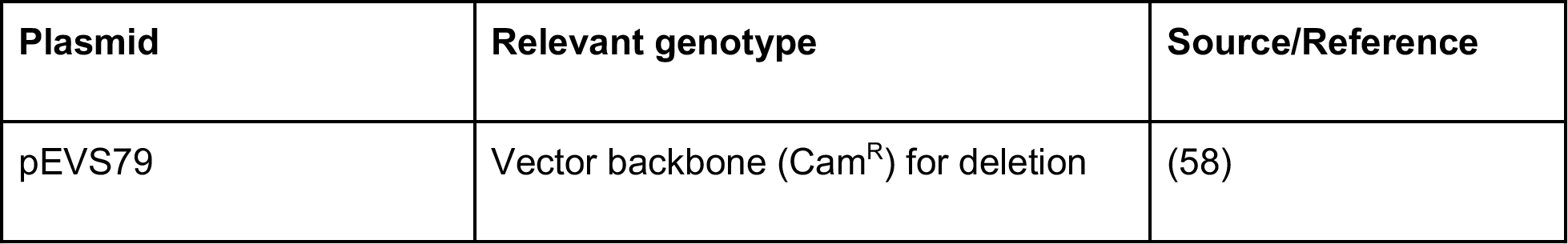
Plasmids.

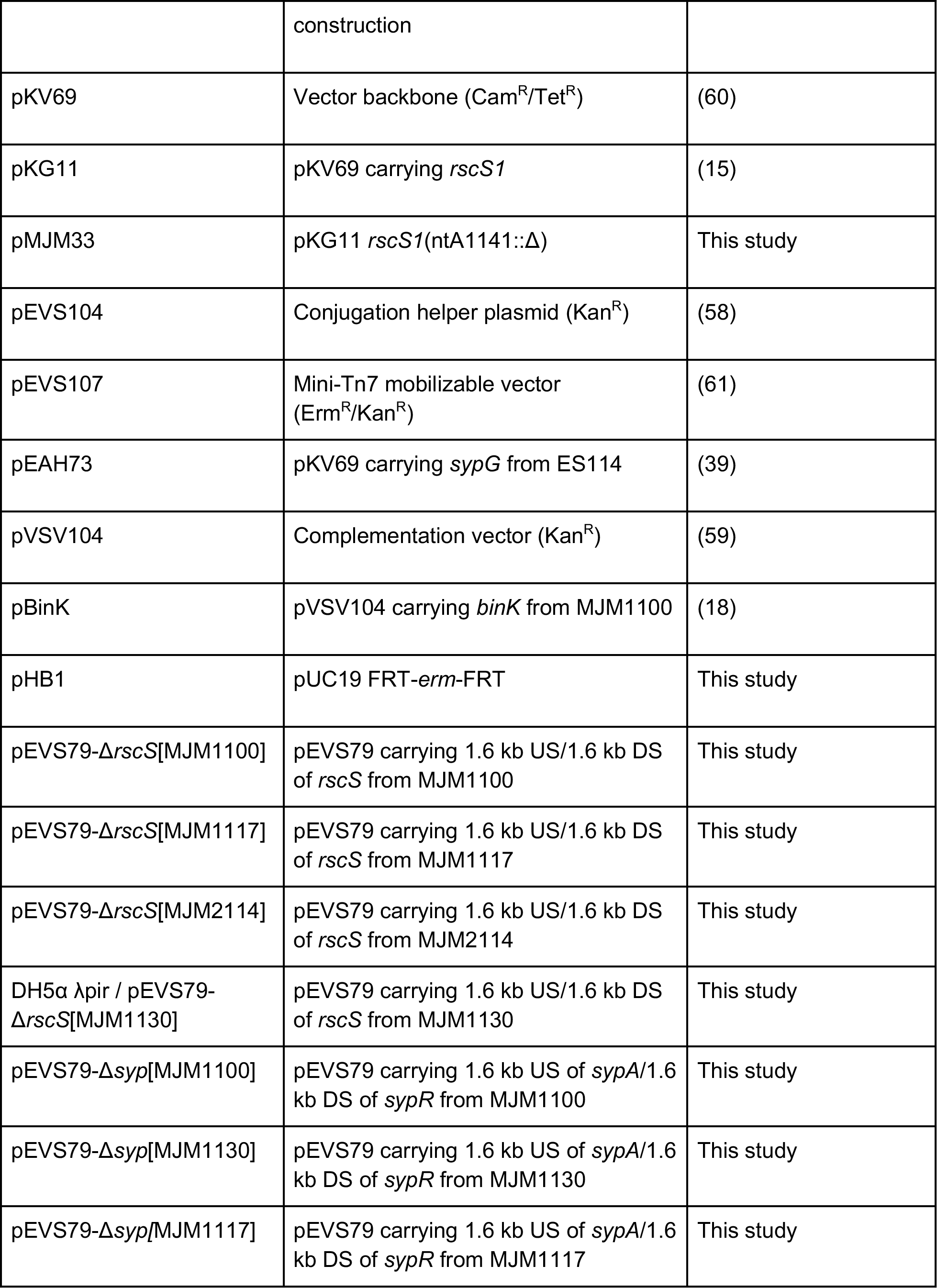

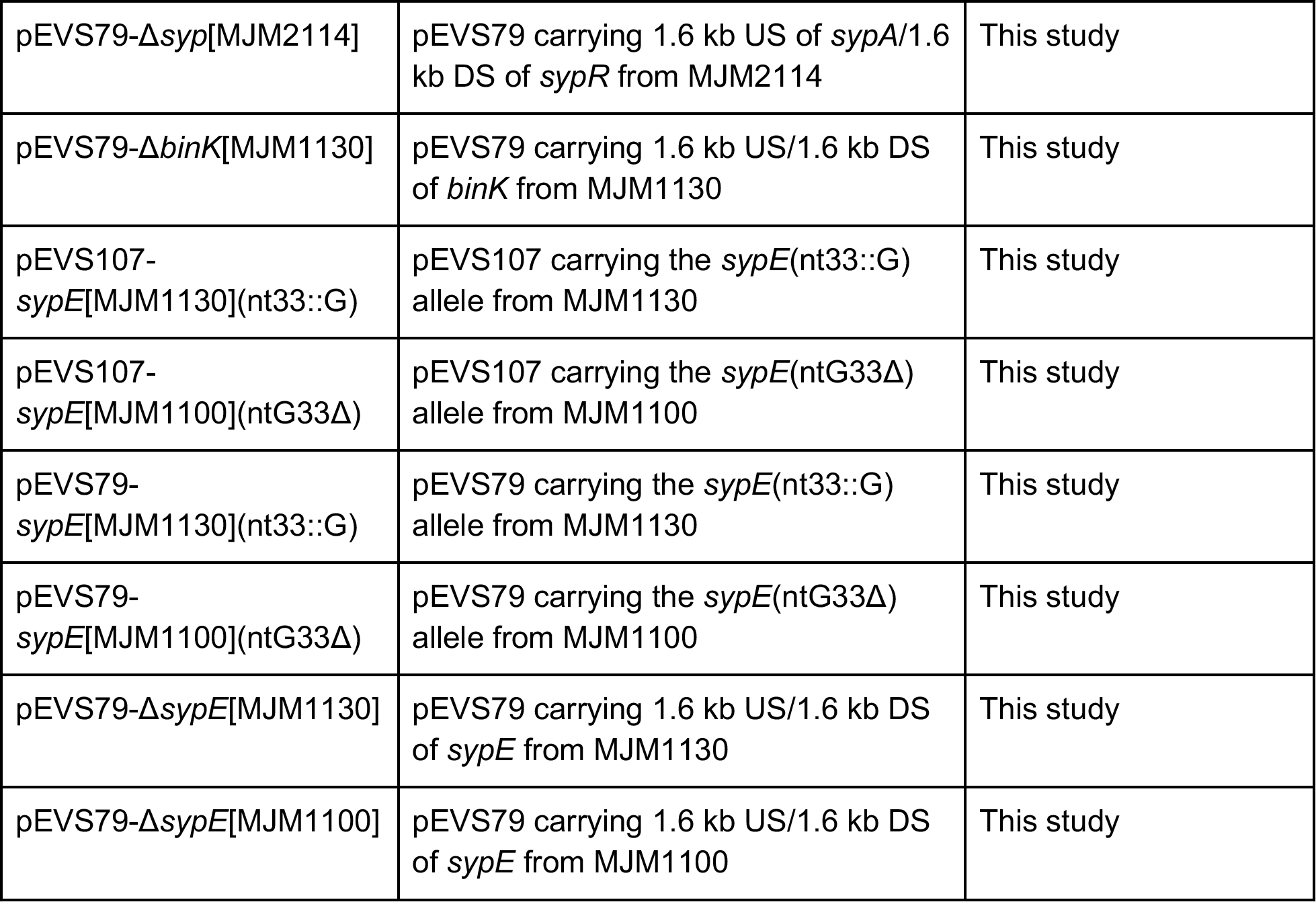

### Phylogenetic analysis

Phylogenetic reconstructions assuming a tree-like topology were created with three methods: maximum parsimony (MP); maximum likelihood (ML); and Bayesian inference (Bayes) as previously described (22, 30). Briefly, MP reconstructions were performed by treating gaps as missing, searching heuristically using random addition, tree-bisection reconnection with a maximum of 8 for swaps, and swapping on best only with 1000 repetitions. For ML and Bayesian analyses, likelihood scores of 1500+ potential evolutionary models were evaluated using both the corrected and uncorrected Akaike Information Criterion, the Bayesian Information Criterion, and Decision Theory (Performance Based Selection) as implemented by jModelTest2.1 (44). For all information criteria, the most optimal evolutionary model was a symmetric model with a proportion of invariable sites and a gamma distribution of rate heterogeneity (SYM+I+Г).

ML reconstruction was implemented via PAUP*4.0a163 (45) by treating gaps as missing, searching heuristically using random addition, tree-bisection reconnection for swaps, and swapping on best only with 1000 repetitions. Bayesian inference was done by invoking the ‘nst=6’ and ‘rates=invgamma’ and ‘statefreqpr=fixed(equal)’ settings in the software package MrBayes3.2.6 (46). The Metropolis-coupled Markov chain Monte Carlo (MCMCMC) algorithm used to estimate the posterior probability distribution for the sequences was set up with ‘temp=0.2’ and one incrementally ‘heated’ chain with three ‘cold’ chains; these four chains were replicated two times per analysis to establish convergence of the Markov chains (i.e., ‘stationarity’ as defined by (47) and interpreted previously in (30)). For this work, stationarity was achieved after approximately 50,000 samples (5,000,000 generations) were collected, with 25% discarded. The ~37,500 samples included were used to construct a 50% majority-rule consensus tree from the sample distribution generated by MCMCMC and assess clades’ posterior probabilities. For ML and MP analyses, the statistical confidence in the topology of each reconstruction was assessed using 1000 bootstrap replicates. Phylogenetic trees were visualized with FigTree 1.4.3 (http://tree.bio.ed.ac.uk/software/figtree); the final tree was edited for publication with Inkscape 0.91 (http://inkscape.org/) and GIMP 2.8.22 (http://www.gimp.org/).

### DNA synthesis and sequencing

Each of the primers listed in Table 3 was synthesized by Integrated DNA Technologies (Coralville, IA). Full inserts from all cloned constructs were verified by Sanger DNA sequencing through ACGT, Inc via the Northwestern University Feinberg School of Medicine NUSeq Core Facility; or the University of Wisconsin-Madison Biotechnology Center. Sequence data was analyzed with SeqMan Pro (DNAStar software), SnapGene (GSL Biotech), and Benchling.

**Table 3.**
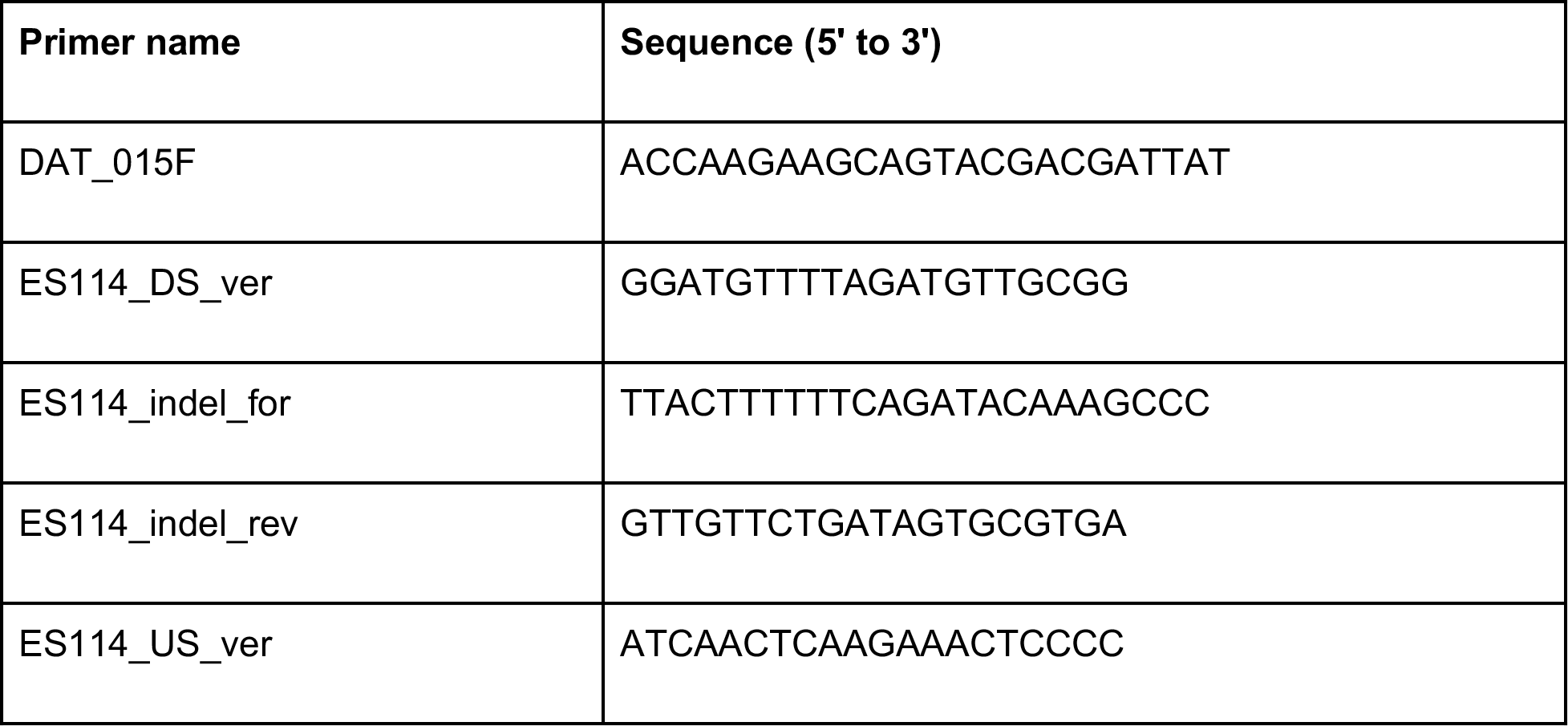
DNA oligonucleotides for PCR amplification and sequencing.

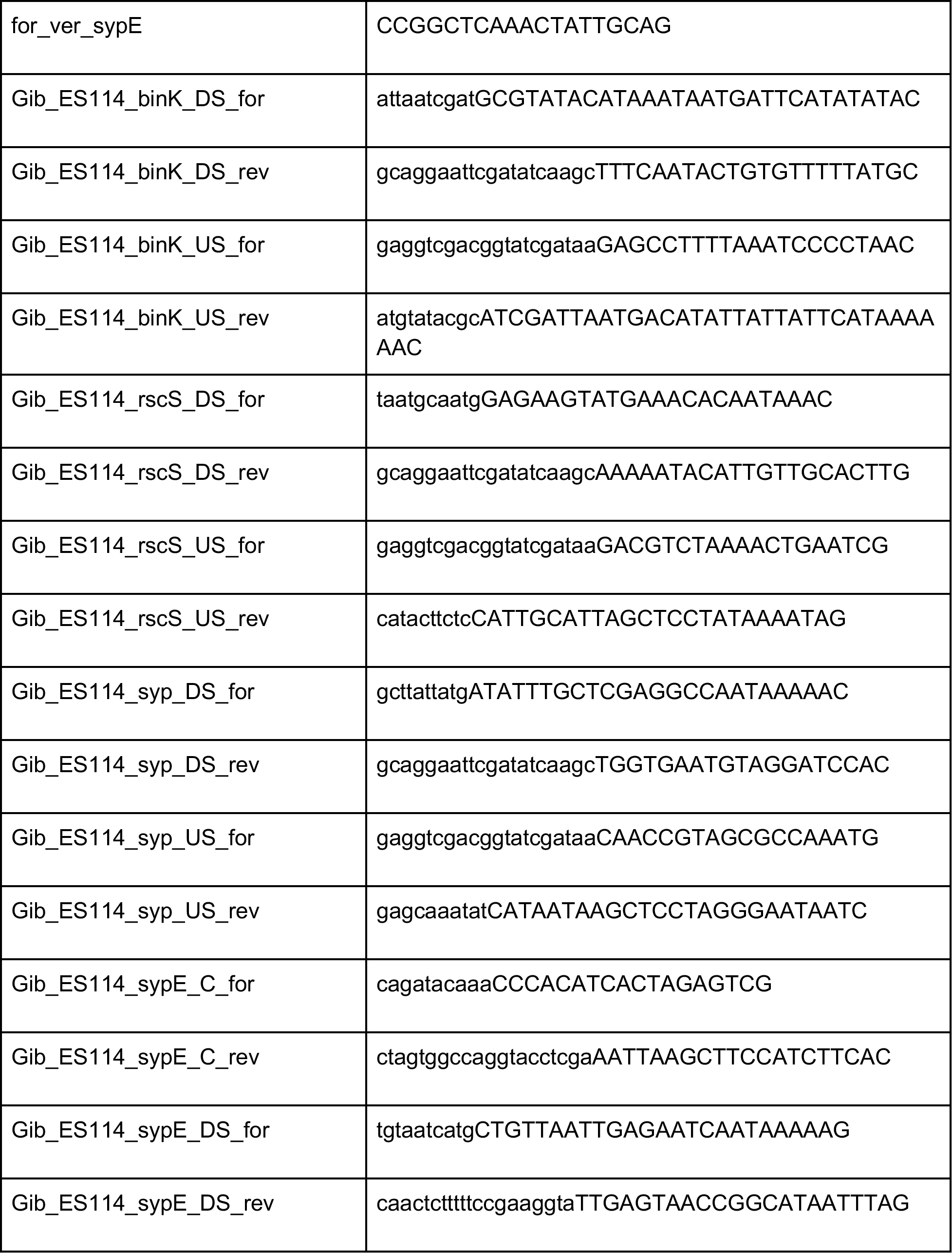

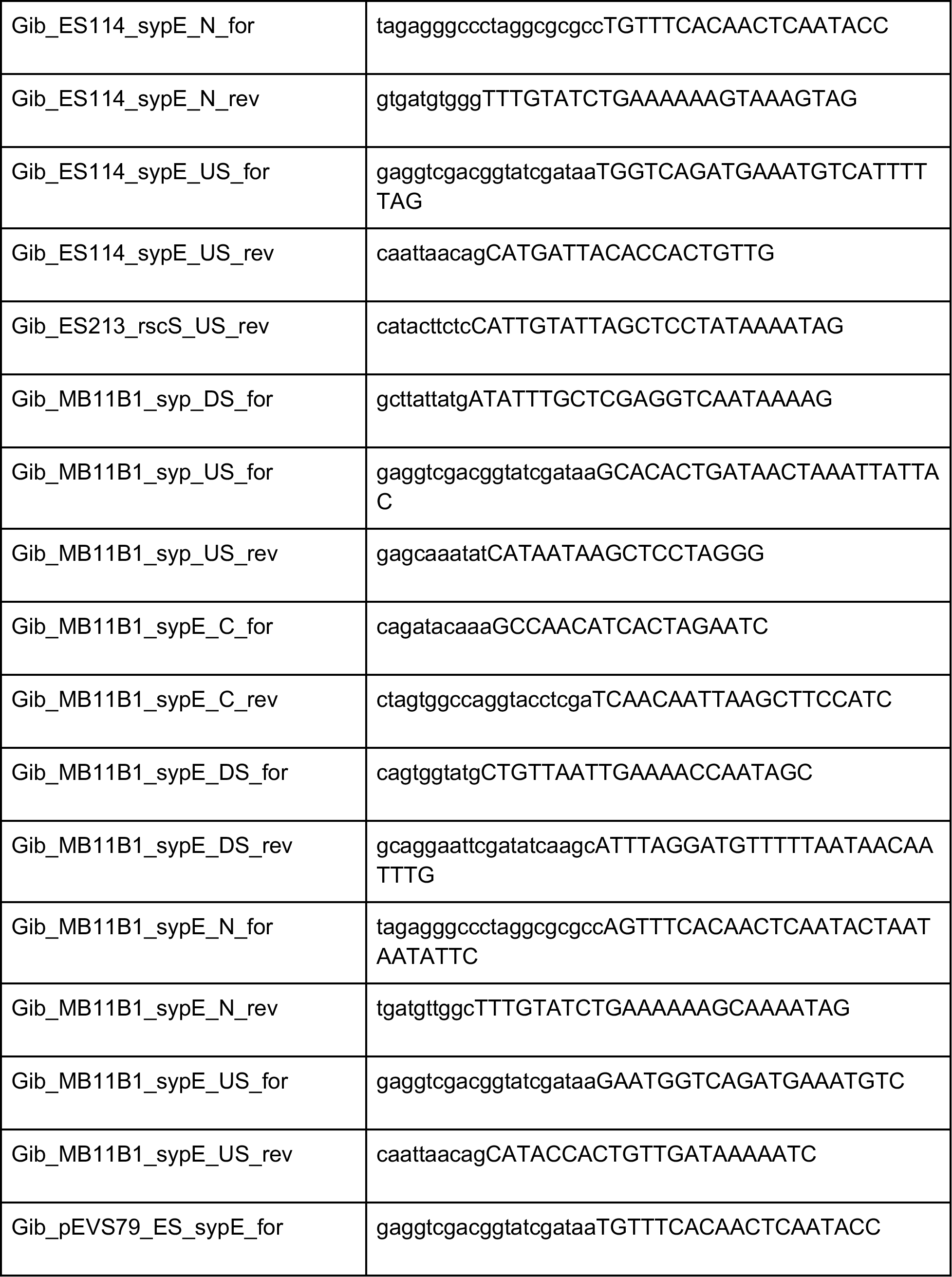

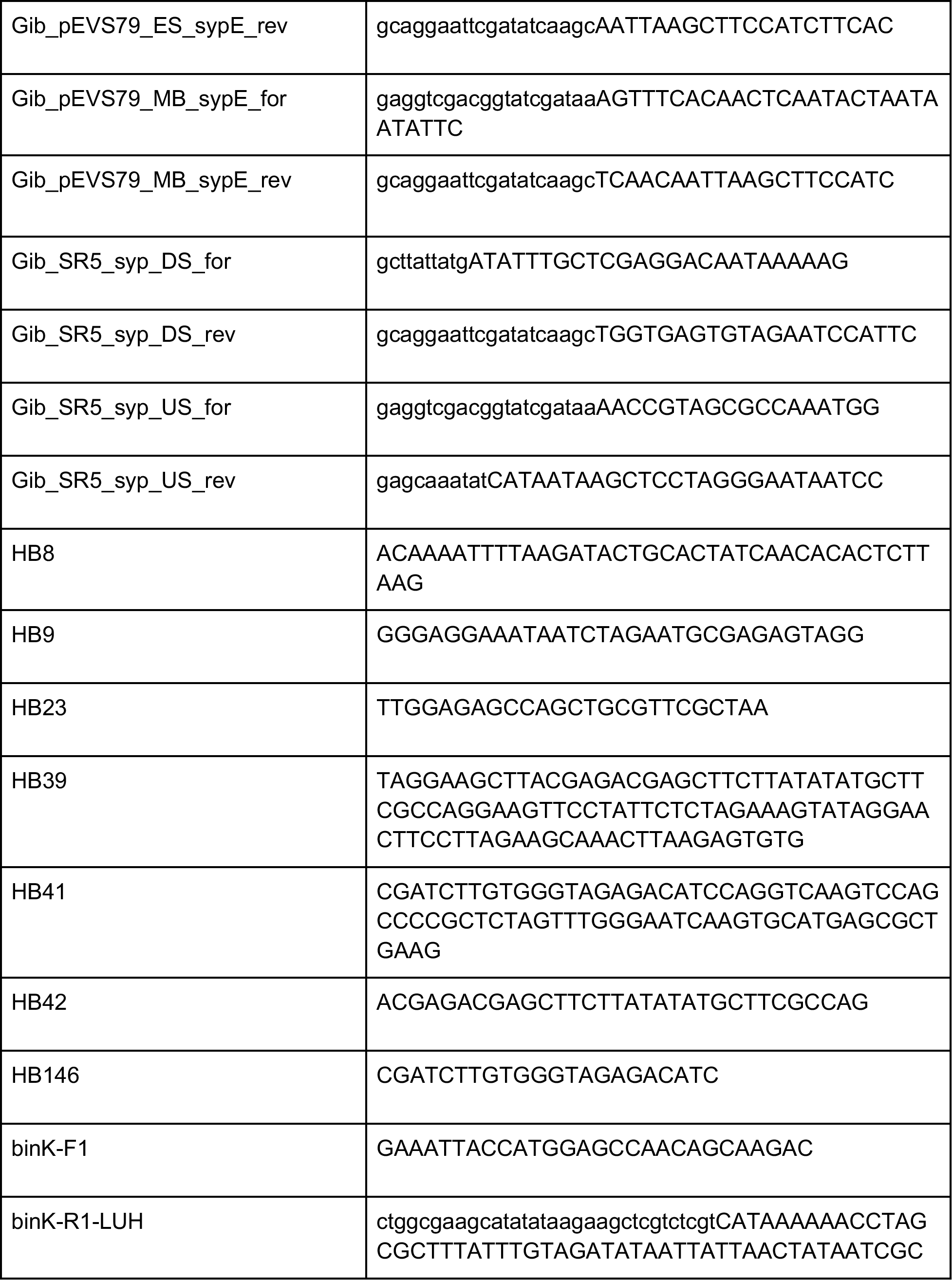

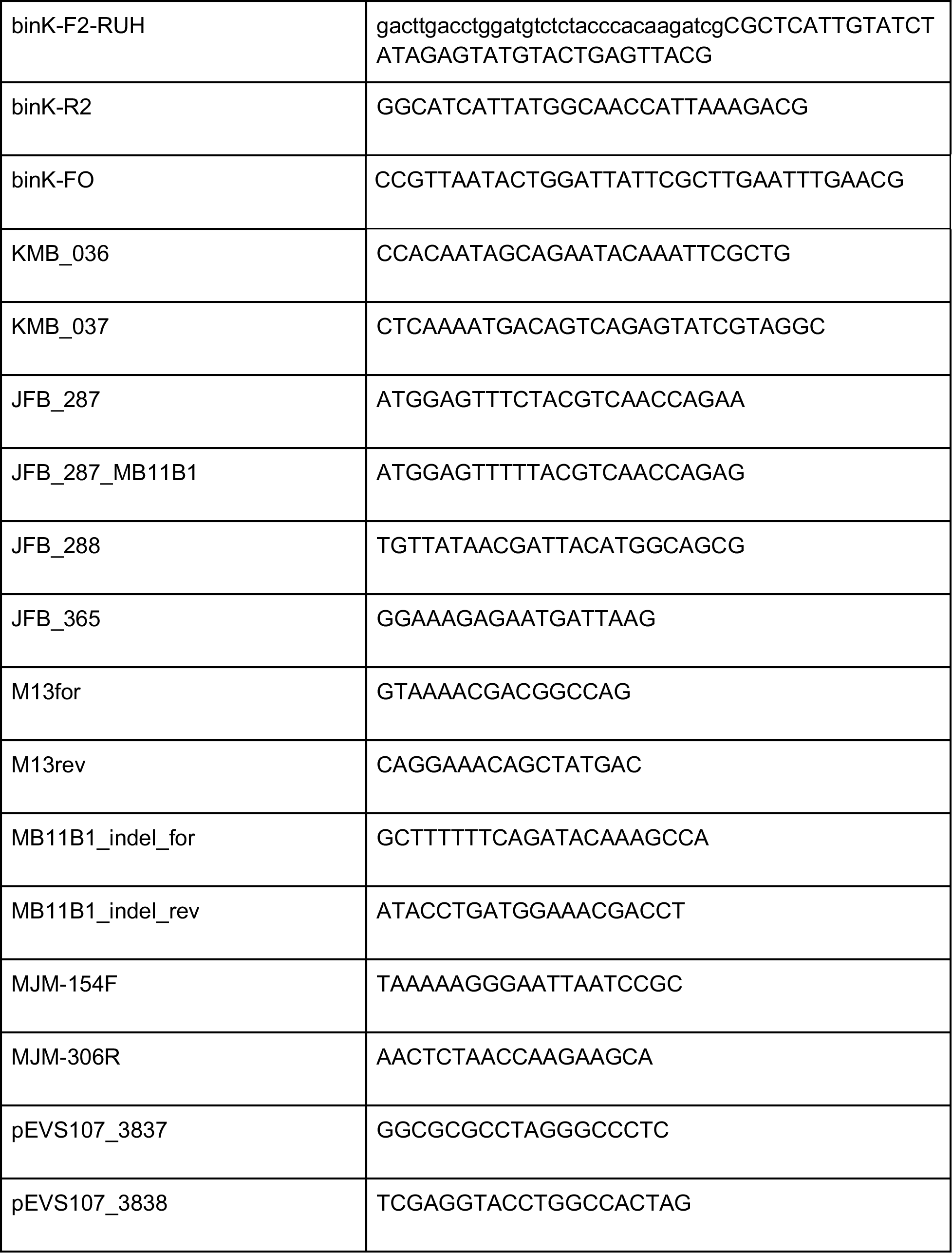

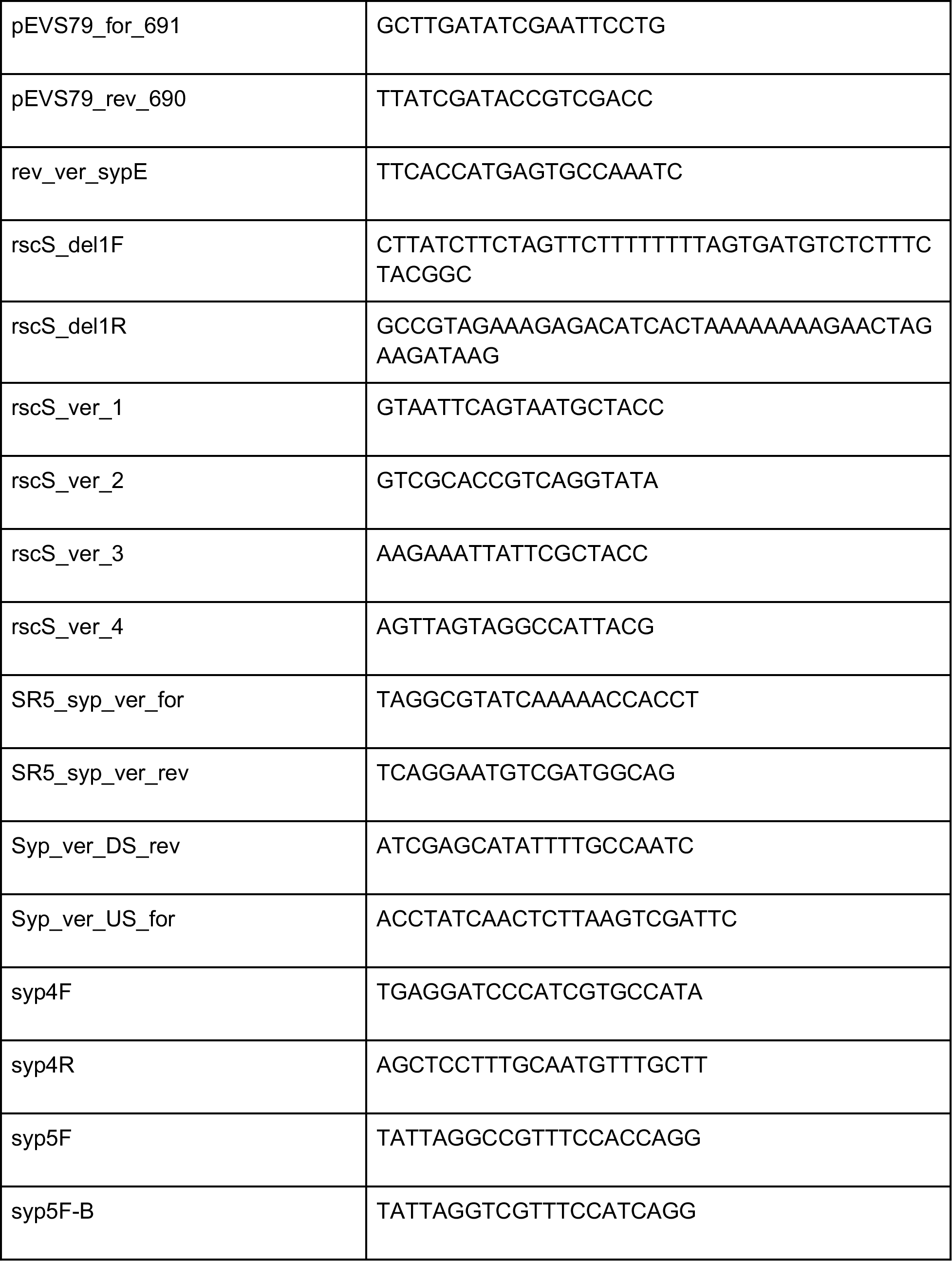

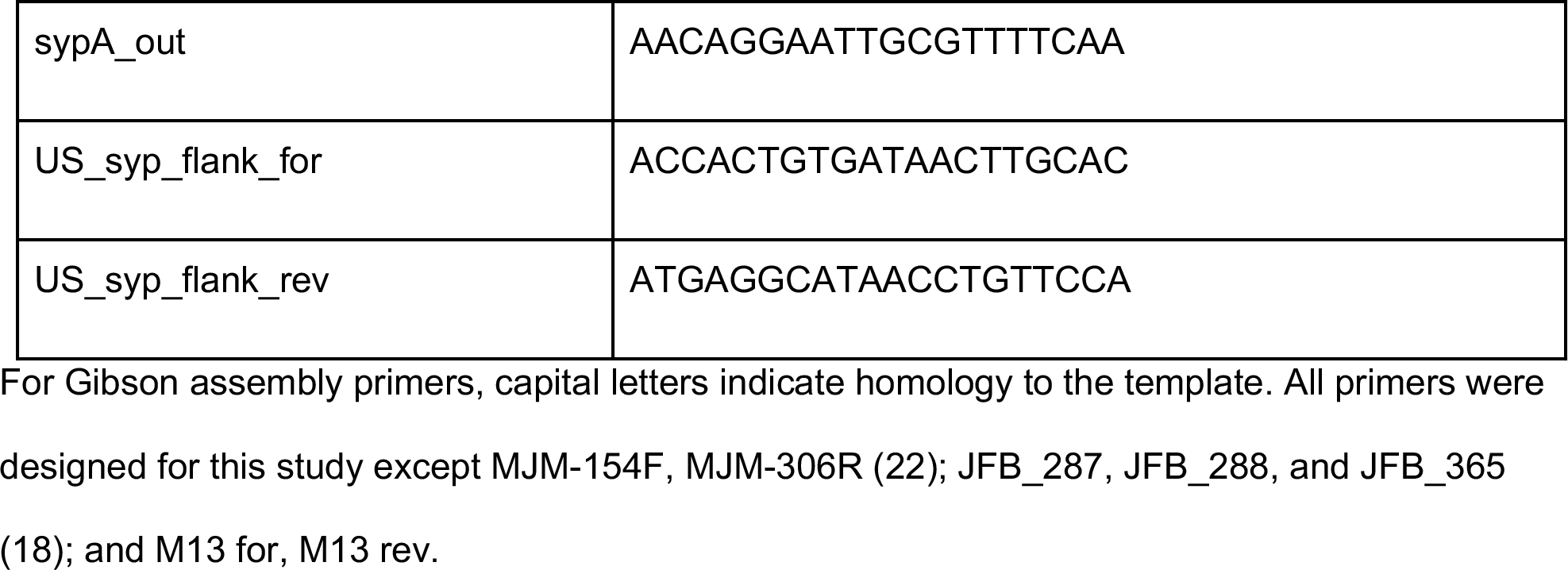

### Construction of gene deletions

Deletions in *V. fischeri* strains ES114 and MB11B1 were made according to the lab’s gene deletion protocol: doi:10.5281/zenodo.1470836. In brief, 1.6 kb upstream and 1.6 kb downstream of the targeted gene or locus were cloned into linearized plasmid pEVS79 (amplified with primers pEVS79_rev_690/pEVS79_for_691) using Gibson Assembly (NEBuilder HiFi DNA Assembly cloning kit) with the primer combinations listed in Table S1. The Gibson mix, linking together the upstream and downstream flanking regions, was transformed into *E. coli* on plates containing X-gal, with several white colonies selected for further screening by PCR using primers flanking the upstream/downstream junction (Tables 3 and S1). The resulting plasmid candidate was confirmed by sequencing and conjugated into the *V. fischeri* recipient by tri-parental mating with helper plasmid pEVS104, selecting for the chloramphenicol resistance of the plasmid backbone. *V. fischeri* colonies were first screened for single recombination into the chromosome by maintaining antibiotic resistance in the absence of selection and then screened for double recombination by the loss of both the antibiotic resistant cassette and the gene/locus of interest. Constructs were verified by PCR (Table 3) and sequencing.

Deletion of SR5 *binK* was conducted using Splicing by Overlap Extension PCR (SOE-PCR) and natural transformation (method modified from (48)). Oligos binK-F1 and binK-R1-LUH, and oligos binK-F2-RUH and binK-R2 were used in a PCR with MJM1125 (SR5) genomic DNA as the template to amplify DNA fragments containing ~1 kb of sequence upstream and downstream relative to *binK*, respectively. Using SOE-PCR, these fragments were fused on either side to a third DNA fragment containing an Erm^R^ cassette, which was amplified using pHB1 as template and oligos HB41 and HB42. We then used natural transformation with pLostfoX (49) to insert this mutagenic DNA into MJM1125, where the flanking sequences guide the Erm^R^ cassette to replace *binK*, generating the desired gene deletion. Candidate SR5 Δ*binK* mutants were selected after growth on LBS-Erm5 plates. Oligos binK-F1 and binK-R2, and HB8 and binK-FO were used to screen candidates for the correct deletion scar by PCR, and oligos KMB_036 and KMB_037 were used to confirm the absence of *binK* in the genome. The deletion was verified by Sanger sequencing with primers HB8, HB9, HB42, and HB146. The base plasmid pHB1 contains an erythromycin resistance cassette flanked by FRT sites, and was constructed using oligos HB23 and HB39 with gBlock gHB1 (sequence in Supplementary File S1; Integrated DNA Technologies, Inc.) as template to amplify the Erm^R^ cassette flanked by HindIII and BamHI sites, which was then cloned into the corresponding site in pUC19.

For most constructs, the deleted genetic material was between the start codon and last six amino acids (50), with two exceptions: the Δ*sypE* in MJM1130 included the ATG that is two amino acids upstream of the predicted start codon, but not the canonical start codon; and the Δ*binK* alleles in MJM1117, MJM1130, and MJM2114, which were constructed to be equivalent to MJM2251 (Δ*binK* in ES114) (18). The Δ*binK* alleles in these strains include the start codon, the next six codons, two codons resulting from ATCGAT (ClaI site), and the last three codons for a predicted 12 amino acid peptide.

### Construction of *sypE* alleles

To create *sypE*(ntG33Δ) in MJM1100 and *sypE*(nt33::G) in MJM1130, the single point mutation was created by amplifying the gene in two halves, with the N-terminal portion consisting of approximately 300 bp upstream of *sypE* up through nucleotide 33 and the C-terminal portion consisting of nucleotide 33 and the remaining *sypE* gene. The overlap between the two halves contained the single nucleotide polymorphism in the primers that connected them. The altered *sypE* alleles were initially cloned into plasmid pEVS107 (linearized with primers pEVS107_3837/pEVS107_3838) using Gibson Assembly and then the entire altered *sypE* allele was subcloned into pEVS79 with Gibson Assembly (Table S1). After double recombination of the vector into *V. fischeri*, candidate colonies for the altered *sypE* in MJM1100 were screened with primers ES114_indel_for/ES114_indel_rev. The primer set anneals more strongly to the wildtype *sypE* sequence than to *sypE*(ntG33::Δ). Candidates in the MJM1100 background with a fainter PCR band were sequenced and confirmed to have the *sypE*(ntG33::Δ) allele. For MJM1130, the primer set MB11B1_indel_for/MB11B1_indel_rev anneals more strongly to the *sypE*(nt33::G) allele than to the naturally occurring *sypE* allele and candidates in MJM1130 that contained a more robust PCR band were selected for sequencing to be confirmed as being *sypE*(nt33::G).

### Construction of pKG11 *rscS1*(ntA1141::Δ)

Plasmid pKG11 encodes an overexpression allele of RscS, termed *rscS1 (15, 28)*. *rscS* nucleotide A1141 was deleted on the plasmid using the Stratagene Quikchange II Site-Directed Mutagenesis Kit with primers rscS_del1F and rscS_del1R. The resulting plasmid, pMJM33, was sequenced with primers MJM-154F and MJM-306R to confirm the single base pair deletion.

### Squid colonization

Hatchling *E. scolopes* were colonized by exposure to approximately 3 × 10^3^ CFU/ml (ranging from 5.2 × 10^2^ - 1.4 × 10^4^ CFU/ml; as specified in figure legends) of each strain in a total volume of 40 ml of FSIO (filter-sterilized Instant Ocean) for 3 hours. Squid were then transferred to 100 ml of FSIO to stop the inoculation and then transferred to 40 ml FSIO for an additional 45 hours with a water change at 24 hours post inoculation. For Figure 10A, kanamycin was added to the FSIO to keep selective pressure on the plasmid. After 48 hours of colonization, the squid were euthanized and surface sterilized by storage at −80 °C, according to standard practices (51). For determination of CFU per light organ, hatchlings were thawed, homogenized, and 50 μl of homogenate dilutions was plated onto LBS plates. Bacterial colonies from each plate were counted and recorded. Mock treated, uncolonized hatchlings (“apo-symbiotic”) were used to determine the limit of detection in the assay. The competitive index (CI) was calculated from the relative CFU of each sample in the output (light organ) versus the input (inoculum) as follows:

Log_10_ ((Test strain[light organ] / Control strain[light organ]) / (Test strain[inoculum] / Control strain[inoculum])). For competitions of natural isolates, the Group A strain (or its Δ*rscS* derivative) was the test strain and the Group B strain was the control strain. Colony color was used to enumerate colonies from each--white for Group A strains MB11B1 and ES213; yellow for Group B strains ES114 and MB15A4--along with PCR verification of selected colonies. For competition between SR5 and SR5 Δ*binK*, 100 colonies per squid were patched onto LBS-Erm5 and LBS.

### Colony biofilm assays

Bacterial strains were grown in LBS media (Fig. 10C) or LBS-Cam2.5 media (Figs. 2, 8) for approximately 17 hours, then 10 μl (Fig. 2) or 8 μl (Fig. 8, 10C) was spotted onto LBS plates (Fig. 10C) or LBS-Tet5 plates (Figs. 2, 8). Spots were allowed to dry and the plates incubated at 25 °C for 48 hours. Images of the spots were taken at 24 and 48 h post-spotting using a Leica M60 microscope and Leica DFC295 camera. After 48 h of growth, the spots were disrupted using a flat toothpick and imaged similarly.

### Analysis of DNA and protein sequences *in silico*

Amino acid sequences for *V. fischeri* ES114 *syp* genes were obtained from RefSeq accession NC_006841.2. Local TBLASTN queries were performed for each protein against nucleotide databases for the following strains, each of which were derived from the RefSeq cds_from_genomic.fna file: *V. fischeri* SR5 (GCA_000241785.1), *V. fischeri* MB11B1 (GCA_001640385.1) and *V. vulnificus* ATCC27562 (GCA_002224265.1). Percent amino acid identity was calculated as the identity in the BLAST query divided by the length of the amino acid sequence in ES114. Domain information is from the PFAM database (52).

## SUPPLEMENTAL FILES

**Table S1.**
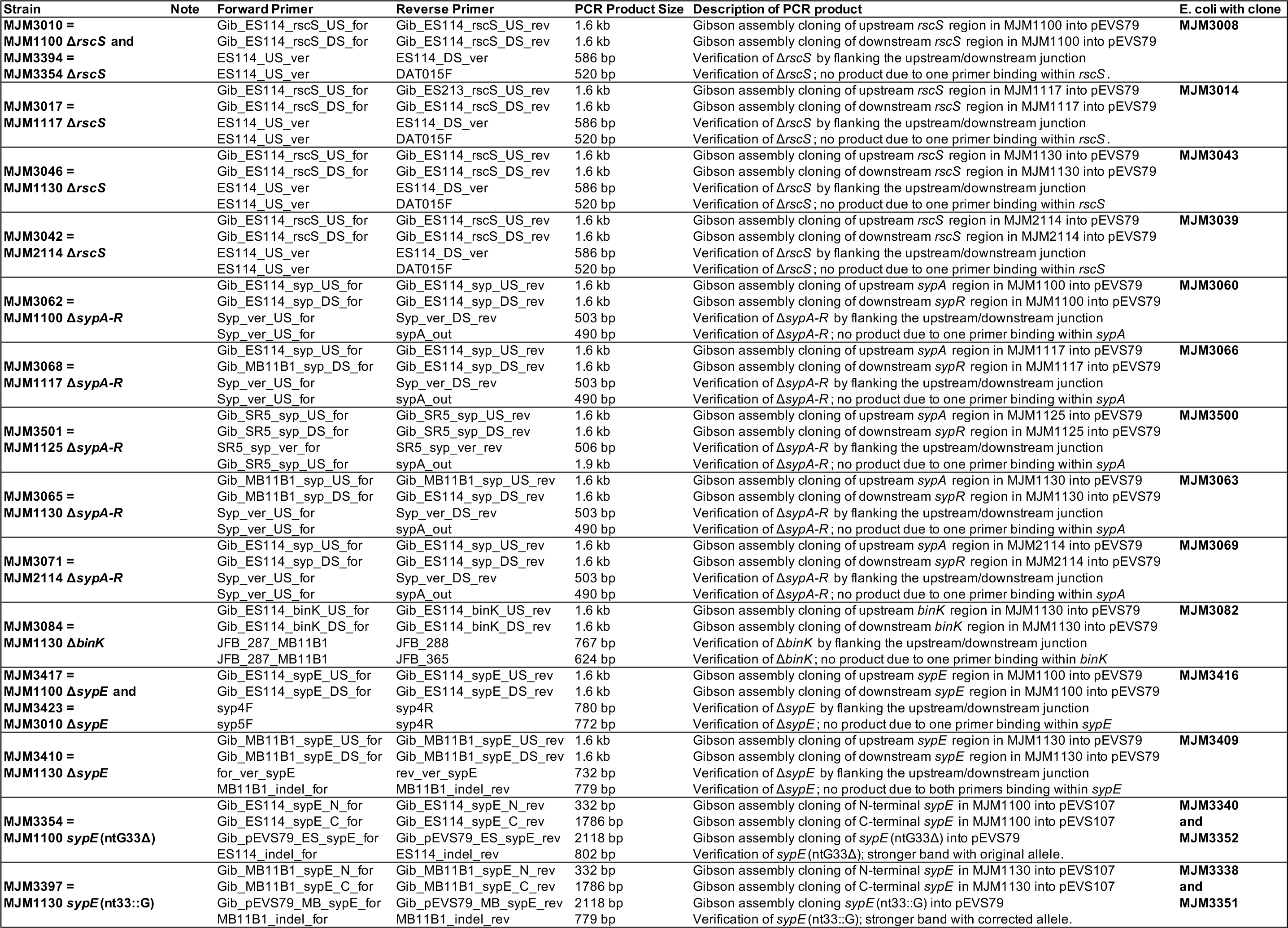
Primer pairs for construction of the deletion mutants. Detailed oligonucleotide and construction details for deletions and the MJM1100 sypE(ntG33Δ) and MJM1130 *sypE*(*nt33*::*G*) strains.

**File S1 (PDF). Sequence of the synthetic dsDNA, gBlock_erm.** The sequence is provided in FASTA format printed as PDF.

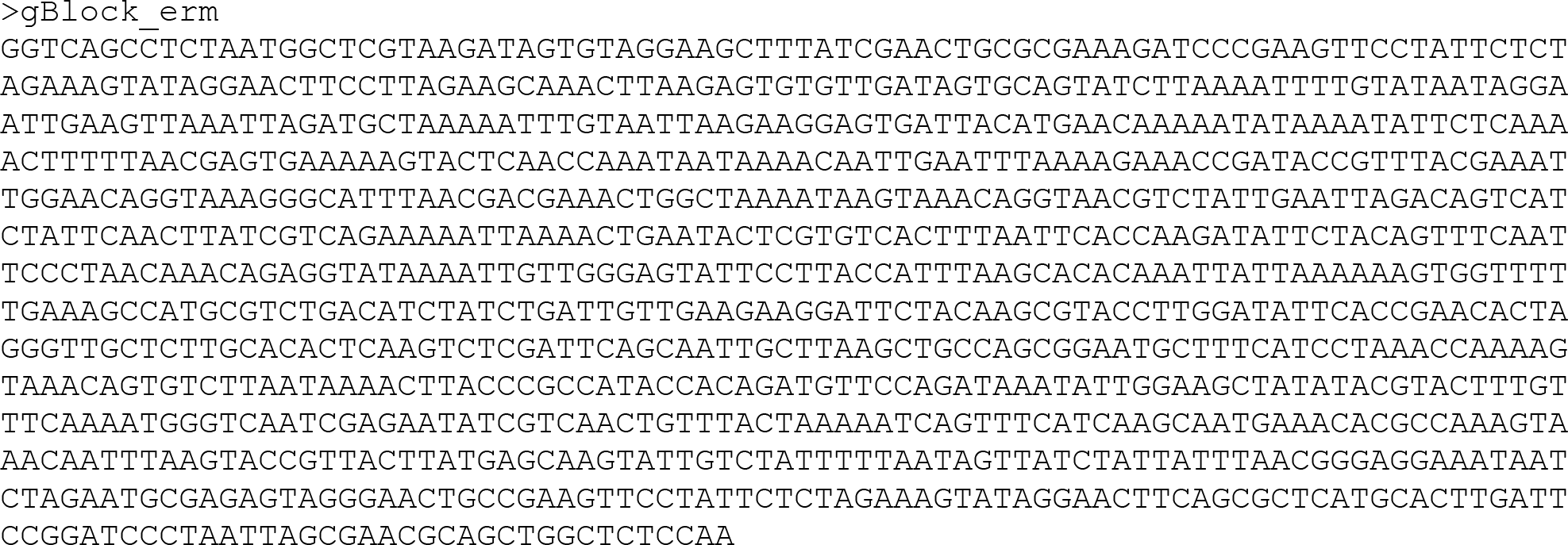

## ACKNOWLEDGMENTS

The authors thank Elizabeth Bacon, Jacklyn Duple, Cheeneng Moua, Lynn Naughton, Olivia Sauls, and Denise Tarnowski for assistance with experiments.

## REFERENCES

1. Long SR. 1996. Rhizobium symbiosis: nod factors in perspective. Plant Cell 8:1885–1898.

2. Roche P, Maillet F, Plazanet C, Debellé F, Ferro M, Truchet G, Promé JC, Dénarié J. 1996. The common nodABC genes of Rhizobium meliloti are host-range determinants. Proc Natl Acad Sci U S A 93:15305–15310.

3. Costello EK, Lauber CL, Hamady M, Fierer N, Gordon JI, Knight R. 2009. Bacterial community variation in human body habitats across space and time. Science 326:1694–1697.

4. Grice EA, Kong HH, Conlan S, Deming CB, Davis J, Young AC, NISC Comparative Sequencing Program, Bouffard GG, Blakesley RW, Murray PR, Green ED, Turner ML, Segre JA. 2009. Topographical and temporal diversity of the human skin microbiome. Science 324:1190–1192.

5. Mark Welch JL, Rossetti BJ, Rieken CW, Dewhirst FE, Borisy GG. 2016. Biogeography of a human oral microbiome at the micron scale. Proc Natl Acad Sci U S A 113:E791–E800.

6. Ruby EG. 2008. Symbiotic conversations are revealed under genetic interrogation. Nat Rev Microbiol 6:752–762.

7. Nyholm SV, McFall-Ngai MJ. 2004. The winnowing: establishing the squid-vibrio symbiosis. Nat Rev Microbiol 2:632–642.

8. Visick KL, Ruby EG. 2006. Vibrio fischeri and its host: it takes two to tango. Curr Opin Microbiol 9:632–638.

9. Mandel MJ, Schaefer AL, Brennan CA, Heath-Heckman EAC, Deloney-Marino CR, McFall-Ngai MJ, Ruby EG. 2012. Squid-derived chitin oligosaccharides are a chemotactic signal during colonization by Vibrio fischeri. Appl Environ Microbiol 78:4620–4626.

10. Jones BW, Nishiguchi MK. 2004. Counterillumination in the Hawaiian bobtail squid, Euprymna scolopes Berry (Mollusca: Cephalopoda). Mar Biol 144:1151–1155.

11. Ruby EG, McFall-Ngai MJ. 1992. A squid that glows in the night: development of an animal-bacterial mutualism. J Bacteriol 174:4865–4870.

12. Wier AM, Nyholm SV, Mandel MJ, Massengo-Tiassé RP, Schaefer AL, Koroleva I, Splinter-Bondurant S, Brown B, Manzella L, Snir E, Almabrazi H, Scheetz TE, Bonaldo M de F, Casavant TL, Soares MB, Cronan JE, Reed JL, Ruby EG, McFall-Ngai MJ. 2010. Transcriptional patterns in both host and bacterium underlie a daily rhythm of anatomical and metabolic change in a beneficial symbiosis. Proc Natl Acad Sci U S A 107:2259–2264.

13. Visick KL. 2009. An intricate network of regulators controls biofilm formation and colonization by Vibrio fischeri. Mol Microbiol 74:782–789.

14. Yip ES, Grublesky BT, Hussa EA, Visick KL. 2005. A novel, conserved cluster of genes promotes symbiotic colonization and σ54-dependent biofilm formation by Vibrio fischeri. Mol Microbiol 57:1485–1498.

15. Yip ES, Geszvain K, DeLoney-Marino CR, Visick KL. 2006. The symbiosis regulator RscS controls the syp gene locus, biofilm formation and symbiotic aggregation by Vibrio fischeri. Mol Microbiol 62:1586–1600.

16. Shibata S, Yip ES, Quirke KP, Ondrey JM, Visick KL. 2012. Roles of the Structural Symbiosis Polysaccharide (syp) Genes in Host Colonization, Biofilm Formation, and Polysaccharide Biosynthesis in Vibrio fischeri. J Bacteriol 194:6736–6747.

17. Morris AR, Visick KL. 2013. The response regulator SypE controls biofilm formation and colonization through phosphorylation of the syp-encoded regulator SypA in Vibrio fischeri. Mol Microbiol 87:509–525.

18. Brooks JF 2nd, Mandel MJ. 2016. The Histidine Kinase BinK Is a Negative Regulator of Biofilm Formation and Squid Colonization. J Bacteriol 198:2596–2607.

19. Tischler AH, Lie L, Thompson CM, Visick KL. 2018. Discovery of Calcium as a Biofilm-Promoting Signal for Vibrio fischeri Reveals New Phenotypes and Underlying Regulatory Complexity. J Bacteriol 200:e00016–18.

20. Pankey MS, Foxall RL, Ster IM, Perry LA, Schuster BM, Donner RA, Coyle M, Cooper VS, Whistler CA. 2017. Host-selected mutations converging on a global regulator drive an adaptive leap towards symbiosis in bacteria. Elife 6:e24414.

21. Thompson CM, Tischler AH, Tarnowski DA, Mandel MJ, Visick KL. 2018. Nitric oxide inhibits biofilm formation by Vibrio fischeri via the nitric oxide-sensor HnoX. Mol Microbiol doi:10.1111/mmi.14147.

22. Mandel MJ, Wollenberg MS, Stabb EV, Visick KL, Ruby EG. 2009. A single regulatory gene is sufficient to alter bacterial host range. Nature 458:215–218.

23. Nyholm SV, Nishiguchi MK. 2008. The evolutionary ecology of a sepiolid squid-vibrio association: from cell to environment. Vie Milieu Paris 58:175–184.

24. Fidopiastis PM, von Boletzky S, Ruby EG. 1998. A new niche for Vibrio logei, the predominant light organ symbiont of squids in the genus Sepiola. J Bacteriol 180:59–64.

25. Ruby EG, Lee KH. 1998. The Vibrio fischeri-Euprymna scolopes Light Organ Association: Current Ecological Paradigms. Appl Environ Microbiol 64:805–812.

26. Mandel MJ. 2010. Models and approaches to dissect host-symbiont specificity. Trends Microbiol 18:504–511.

27. Gyllborg MC, Sahl JW, Cronin DC 3rd, Rasko DA, Mandel MJ. 2012. Draft genome sequence of Vibrio fischeri SR5, a strain isolated from the light organ of the Mediterranean squid Sepiola robusta. J Bacteriol 194:1639.

28. Geszvain K, Visick KL. 2008. Multiple factors contribute to keeping levels of the symbiosis regulator RscS low. FEMS Microbiol Lett 285:33–39.

29. Wollenberg MS, Ruby EG. 2009. Population structure of Vibrio fischeri within the light organs of Euprymna scolopes squid from Two Oahu (Hawaii) populations. Appl Environ Microbiol 75:193–202.

30. Wollenberg MS, Ruby EG. 2012. Phylogeny and fitness of Vibrio fischeri from the light organs of Euprymna scolopes in two Oahu, Hawaii populations. ISME J 6:352–362.

31. Elliott KT, DiRita VJ. 2008. Characterization of CetA and CetB, a bipartite energy taxis system in Campylobacter jejuni. Mol Microbiol 69:1091–1103.

32. Antonov I, Coakley A, Atkins JF, Baranov PV, Borodovsky M. 2013. Identification of the nature of reading frame transitions observed in prokaryotic genomes. Nucleic Acids Res 41:6514–6530.

33. Bongrand C, Koch EJ, Moriano-Gutierrez S, Cordero OX, McFall-Ngai M, Polz MF, Ruby EG. 2016. A genomic comparison of 13 symbiotic Vibrio fischeri isolates from the perspective of their host source and colonization behavior. ISME J 10:2907–2917.

34. Altschul SF, Madden TL, Schäffer AA, Zhang J, Zhang Z, Miller W, Lipman DJ. 1997. Gapped BLAST and PSI-BLAST: a new generation of protein database search programs. Nucleic Acids Res 25:3389–3402.

35. Rusch DB, Rowe-Magnus DA. 2017. Complete Genome Sequence of the Pathogenic Vibrio vulnificus Type Strain ATCC 27562. Genome Announc 5:e00907–17.

36. Morris AR, Darnell CL, Visick KL. 2011. Inactivation of a novel response regulator is necessary for biofilm formation and host colonization by Vibrio fischeri. Mol Microbiol 82:114–130.

37. Guo Y, Rowe-Magnus DA. 2011. Overlapping and unique contributions of two conserved polysaccharide loci in governing distinct survival phenotypes in Vibrio vulnificus. Environ Microbiol 13:2888–2990.

38. Morris AR, Visick KL. 2013. Inhibition of SypG-induced biofilms and host colonization by the negative regulator SypE in Vibrio fischeri. PLoS One 8:e60076.

39. Hussa EA, Darnell CL, Visick KL. 2008. RscS functions upstream of SypG to control the syp locus and biofilm formation in Vibrio fischeri. J Bacteriol 190:4576–4583.

40. Koehler S, Gaedeke R, Thompson C, Bongrand C, Visick KL, Ruby E, McFall-Ngai M. 2018. The model squid-vibrio symbiosis provides a window into the impact of strain- and species-level differences during the initial stages of symbiont engagement. Environ Microbiol doi:10.1111/1462-2920.14392.

41. Ray VA, Visick KL. 2012. LuxU connects quorum sensing to biofilm formation in Vibrio fischeri. Mol Microbiol 86:954–970.

42. Giraud E, Moulin L, Vallenet D, Barbe V, Cytryn E, Avarre J-C, Jaubert M, Simon D, Cartieaux F, Prin Y, Bena G, Hannibal L, Fardoux J, Kojadinovic M, Vuillet L, Lajus A, Cruveiller S, Rouy Z, Mangenot S, Segurens B, Dossat C, Franck WL, Chang W-S, Saunders E, Bruce D, Richardson P, Normand P, Dreyfus B, Pignol D, Stacey G, Emerich D, Verméglio A, Médigue C, Sadowsky M. 2007. Legumes symbioses: absence of Nod genes in photosynthetic bradyrhizobia. Science 316:1307–1312.

43. Bonaldi K, Gargani D, Prin Y, Fardoux J, Gully D, Nouwen N, Goormachtig S, Giraud E. 2011. Nodulation of Aeschynomene afraspera and A. indica by photosynthetic Bradyrhizobium Sp. strain ORS285: the Nod-dependent versus the Nod-independent symbiotic interaction. Mol Plant Microbe Interact 24:1359–1371.

44. Darriba D, Taboada GL, Doallo R, Posada D. 2012. jModelTest 2: more models, new heuristics and parallel computing. Nat Methods 9:772.

45. Swofford DL. 2003. PAUP*: Phylogenetic Analysis Using Parsimony (* and Other Methods) 4th edn. Sinauer.

46. Ronquist F, Teslenko M, van der Mark P, Ayres DL, Darling A, Höhna S, Larget B, Liu L, Suchard MA, Huelsenbeck JP. 2012. MrBayes 3.2: efficient Bayesian phylogenetic inference and model choice across a large model space. Syst Biol 61:539–542.

47. Ronquist F, van der Mark P, Huelsenbeck JP. 2009. Bayesian phylogenetic analysis using MrBayes, 2nd edn. Cambridge University Press.

48. Visick KL, Hodge-Hanson KM, Tischler AH, Bennett AK, Mastrodomenico V. 2018. Tools for Rapid Genetic Engineering of Vibrio fischeri. Appl Environ Microbiol 84:e00850–18.

49. Pollack-Berti A, Wollenberg MS, Ruby EG. 2010. Natural transformation of Vibrio fischeri requires tfoX and tfoY. Environ Microbiol 12:2302–2311.

50. Baba T, Ara T, Hasegawa M, Takai Y, Okumura Y, Baba M, Datsenko KA, Tomita M, Wanner BL, Mori H. 2006. Construction of Escherichia coli K-12 in-frame, single-gene knockout mutants: the Keio collection. Mol Syst Biol 2:2006.0008.

51. Naughton LM, Mandel MJ. 2012. Colonization of Euprymna scolopes squid by Vibrio fischeri. J Vis Exp e3758.

52. Finn RD, Coggill P, Eberhardt RY, Eddy SR, Mistry J, Mitchell AL, Potter SC, Punta M, Qureshi M, Sangrador-Vegas A, Salazar GA, Tate J, Bateman A. 2016. The Pfam protein families database: towards a more sustainable future. Nucleic Acids Res 44:D279–85.

53. Lee K-H. 1994. Ecology of Vibrio fisheri: the light organ symbiont of the Hawaiian sepiolid squid Euprymna scolopes. University of Southern California.

54. Boettcher KJ, Ruby EG. 1990. Depressed light emission by symbiotic Vibrio fischeri of the sepiolid squid Euprymna scolopes. J Bacteriol 172:3701–3706.

55. Boettcher KJ, Ruby EG. 1994. Occurrence of plasmid DNA in the sepiolid squid symbiont Vibrio fischeri. Curr Microbiol 29:279–286.

56. Nishiguchi MK, Ruby EG, McFall-Ngai MJ. 1998. Competitive dominance among strains of luminous bacteria provides an unusual form of evidence for parallel evolution in Sepiolid squid-vibrio symbioses. Appl Environ Microbiol 64:3209–3213.

57. Nishiguchi MK, Nair VS. 2003. Evolution of symbiosis in the Vibrionaceae: a combined approach using molecules and physiology. Int J Syst Evol Microbiol 53:2019–2026.

58. Stabb EV, Ruby EG. 2002. RP4-based plasmids for conjugation between Escherichia coli and members of the Vibrionaceae. Methods Enzymol 358:413–426.

59. Dunn AK, Millikan DS, Adin DM, Bose JL, Stabb EV. 2006. New rfp- and pES213-derived tools for analyzing symbiotic Vibrio fischeri reveal patterns of infection and lux expression in situ. Appl Environ Microbiol 72:802–810.

60. Visick KL, Skoufos LM. 2001. Two-component sensor required for normal symbiotic colonization of Euprymna scolopes by Vibrio fischeri. J Bacteriol 183:835–842.

61. McCann J, Stabb EV, Millikan DS, Ruby EG. 2003. Population dynamics of Vibrio fischeri during infection of Euprymna scolopes. Appl Environ Microbiol 69:5928–5934.

